# A complete digital karyotype of the B-cell leukemia REH cell line resolved by long-read sequencing

**DOI:** 10.1101/2023.03.08.531483

**Authors:** Mariya Lysenkova Wiklander, Gustav Arvidsson, Ignas Bunikis, Anders Lundmark, Amanda Raine, Yanara Marincevic-Zuniga, Henrik Gezelius, Anna Bremer, Lars Feuk, Adam Ameur, Jessica Nordlund

## Abstract

The B-cell acute lymphoblastic leukemia (ALL) cell line REH, with the t(12;21) *ETV6-RUNX1* translocation, is known to have a complex karyotype defined by a series of large-scale chromosomal rearrangements. Taken from a 15-year-old at relapse, the cell line offers a practical model for the study of high-risk pediatric B-ALL patients. In recent years, short-read DNA and RNA sequencing have emerged as a complement to analog karyotyping techniques in the resolution of structural variants in an oncological context. However, it is challenging to create a comprehensive digital karyotype of a genome with these techniques alone. Here, we explore the integration of long-read PacBio and Oxford Nanopore whole genome sequencing (WGS), IsoSeq RNA-sequencing, and short-read sequencing to create a detailed digital karyotype of the REH cell line. WGS refined the breakpoints of known aberrations and clarified the molecular traits of disrupted ALL-associated genes *BTG1* and *TBL1XR1*, as well as the glucocorticoid receptor *NR3C1*. Several previously underreported structural variants were also uncovered, including deletions affecting the ALL-associated genes *VPREB1* and *NFATC1*. Meanwhile, transcriptome sequencing identified seven fusion genes within the genomic breakpoints. Together, our extensive whole-genome investigation makes high-quality open-source data available to the leukemia genomics community.

**KEY POINTS:**

- A complete digital karyotype of the REH cell line was produced with short- and long-read DNA and RNA sequencing technologies.
- The study enabled precise identification of structural variants, and the fusion genes expressed as the result of these variants.

## INTRODUCTION

REH, established in 1974 as the first B cell precursor acute lymphoblastic leukemia (ALL) cell line ^1^, was derived from the peripheral blood of a 15 year-old female patient at relapse ^2^. REH has a near-diploid karyotype characterized by complex structural rearrangements, including several interchromosomal translocations. Among the features described in this cell line is a complex four-way translocation t(4;12;21;16) that results in the canonical subtype-defining *ETV6-RUNX1* fusion gene ^3^ and glucocorticoid (GC) resistance attributed to lack of a GC receptor ^4^. While this cell line is used extensively in leukemia research ^5–10^, its complex karyotype has never been comprehensively characterized.

Traditional karyotyping and banding techniques can resolve structural variants to a resolution of 5Mbp, while FISH and microarray analyses are limited to approximately 150kbp ^11,12^. Whole genome and transcriptome sequencing (WGTS) have become increasingly important as a comprehensive clinical test for accurately resolving genomic breakpoints of structural variants (SVs) and fusion genes in cancer genomes, providing significant improvements in diagnostic yield over analog methods ^13–15^. Sequencing-based diagnostic approaches are predominantly based on short-read next generation sequencing (NGS) technologies, which have benefits in low cost, high accuracy, and extensively validated computational pipelines ^16–18^. However, inherent challenges exist with short-read sequencing, which include difficulty in mapping short reads that arise from repetitive sequences, sequencing through regions of high GC content, and detecting complex SVs ^19,20^. The “dark” regions of the human genome have been difficult to assemble and are thus poorly covered by standard short-read sequencing, preventing identification of disease-relevant SVs within these regions ^21^. Long-read sequencing technologies, including those developed by Pacific Biosciences (PacBio) and Oxford Nanopore Technologies (ONT), have significantly advanced detection and analysis of the dark regions of the genome with their unique capability to read through repetitive regions with variable GC-content, and ability to resolve single-nucleotide variants (SNVs) and SVs even in the most challenging genomic regions ^16,22,23^. Long reads have shown superiority compared to short reads for SV detection, especially when used in combination with multiple SV-calling softwares ^24^. Importantly, long-read technologies have recently made remarkable improvements in throughput and base calling accuracy, which is now over 99% ^25^, making them an interesting alternative for future clinical cancer diagnostics.

There remain numerous clinically or diagnostically important genomic regions overlooked by short-read NGS that have the potential to be resolved by long-read technologies. In the present study, we explore the utility of short- and long-read sequencing technologies to produce a complete digital karyotype of the highly complex REH cell line for future leukemia research. Additionally, we compare and contrast the performance of the different sequencing technologies and analytical pipelines used to produce a high resolution reference dataset for this important cell line.

## METHODS

### REH cell line

The human pre B-cell ALL cell line REH was obtained from DSMZ (ACC 22) and cultured according to the supplier’s specifications. DNA and RNA were extracted from 5M cells using QIAGEN’s AllPrep DNA/RNA Micro Kit. For the long-read WGS libraries, Circulomics Nanobind UHMW DNA Extraction was used with an input of 15M cells. Conventional karyotyping (G-banding) was performed by The Department of Clinical Genetics, Uppsala University Hospital, Uppsala, Sweden (see Supplemental Methods).

### Data Set

We generated three WGS and three RNA-seq datasets from REH cells (**Table 1**). An Illumina PCR-free WGS library was generated with 1000 ng input DNA and sequenced PE-150 on a HiSeq X instrument. An Illumina short-read RNA-seq library was generated with 300 ng input total RNA and the TruSeq stranded total RNA kit, and sequenced PE-100 on a NovaSeq 6000 instrument. One Oxford Nanopore WGS library was generated from 40 μg input DNA prepared with the Ultra-Long Sequencing Kit (SQK-ULK001) and sequenced on the PromethION 24. Two PacBio IsoSeq RNA-seq datasets and one PacBio SMRTbell HiFi WGS dataset were prepared with the SMRTbell^®^ Express Template Prep Kit 2.0, Sequel^®^ II Binding Kit 2.0, and Sequel^®^ II Sequencing Plate 2.0. For the IsoSeq library, we used 300 ng input RNA for each of two libraries; we used a varying bead ratio to generate one library with standard-length transcripts and one library with full-length transcripts. FLNC reads were generated using PacBio tools ccs-5.0.0, lima v2.0.1 and refine v3.4.0. For the PacBio WGS, we used 10 μg input DNA to create a CLR library and 15 μg to create a HiFi library; the resulting CCS reads from both libraries were merged for analysis.

**Table 1.** An overview of the sequencing and analysis methods.

|  |  | Short-read (Illumina) | Long-read (PacBio) | Long-read (ONT) |
| --- | --- | --- | --- | --- |
| WGS | *Library preparation* | TruSeq DNA PCR-Free | SMRTbell | SQK-LSK110 Ligation Sequencing kit |
|  | *Instrument* | HiSeq X Ten | Sequel II | PromethION 24 |
|  | *Raw Reads (GB)* | 72 | 0.7 (CLR)<br>45.2 (HiFi) | 55 |
|  | *Analysis tools* | Sarek, TIDDIT | pbmm2, Sniffles2 | Minimap2, Sniffles2 |
| RNA-seq | *Library preparation* | TruSeq stranded total RNA | SMRTbell IsoSeq |  |
|  | *Instrument* | NovaSeq 6000 | Sequel II |  |
|  | *Raw Reads (GB)* | 4 | 4.9 (standard-length)<br>9.1 (full-length) |  |
|  | *Analysis tools* | nf-core/<br>rnafusion | Minimap2,<br>cDNA_Cupcake,<br>JAFFAL |  |

We also generated one RNA-seq dataset from GM12878 (B lymphoblastoid cell line) by merging four Illumina short-read RNA-seq libraries, each generated using 500 ng of input total RNA with the TruSeq Stranded Total RNA with RiboZero depletion, and sequenced on a NovaSeq 6000 instrument.

### Data Analysis

SV and fusion gene detection were performed by first aligning reads to the reference genome version GRCh38, followed by analysis with SV calling software and fusion gene detection tools, respectively (**Table 1**). For gene annotation, GENCODE V37 (Ensembl 103) was used.

### SV detection using WGS data

For the short-read WGS data, we used Nextflow v21.10.6 ^26^ to run the nf-core/sarek pipeline v2.7 ^27,28^, which handles mapping, indexing, sorting, duplicate marking and recalibration; the pipeline’s TIDDIT module ^29^ was used for SV calling. We mapped and sorted the PacBio CCS reads using pbmm2 v1.9.0 (https://github.com/PacificBiosciences/pbmm2), which uses minimap2 ^30^. After adapter sequences were trimmed using Porechop (v0.2.4, https://github.com/rrwick/Porechop), the ONT reads were mapped using minimap2 v2.24-r1122, followed by sorting and indexing with samtools v1.15.1 ^31^. For both long-read WGS datasets, we ran Sniffles2 v2.0.6 ^32^ for SV detection, with non-germline mode enabled. SURVIVOR 1.0.7 ^33^ was used to filter and merge VCF files using a minimum size of 100bp and excluding variants found in the ENCODE

Blacklist ^34^. For consensus callsets, SVs were required to agree on type and strand, with breakpoints a maximum allowed distance of 1kb. For the evaluation of large-scale variants and translocations, the callsets were programmatically pre-filtered (see Supplemental Methods) and the remaining candidates were inspected manually. SV candidates were confirmed visually in IGV v2.13.0 ^35^, requiring the presence of split long reads and/or discordant mates in at least two of the three SV callsets for confirmation.

### Fusion gene detection

We ran the nf-core/rnafusion pipeline v2.0.0 ^27,36^ on short-read REH RNA-seq data to call and visualize putative fusion genes. This pipeline uses five different fusion tools: Arriba ^37^, FusionCatcher ^38^, pizzly ^39^, squid ^40^ and STAR-fusion ^41^. We also ran nf-core/rnafusion on the GM12878 short-read RNA-seq dataset in order to generate a set of non-leukemic fusion gene candidates for filtering.

Putative fusion genes were called in the long-read IsoSeq data with two different workflows. First, we used cDNA_Cupcake v28.0.0 (https://github.com/Magdoll/cDNA_Cupcake) in conjunction with minimap2, clustering using isoseq3 v3.7.0, and fusion gene classification using SQANTI3 v5.0 ^42^. Second, we used the JAFFAL pipeline v2.1 ^43^, built from the provided Dockerfile and run with default settings.

Candidate fusion genes were programmatically pre-filtered (see Supplemental Methods) and the remaining candidates were inspected manually in IGV.

### Data Sharing Statement

Sequencing data and BAM files are available at NCBI’s Sequence Read Archive (SRA) under BioProject accessions PRJNA600820 and PRJNA834955 (**Supplemental Table S1**). Supplemental data are available at https://doi.org/10.5281/zenodo.7702098. Analysis scripts are available at https://github.com/Molmed/REH.

## RESULTS

### Karyotyping

We used G-banding to confirm the integrity of the REH cells used for sequencing, and to verify the presence of the chromosomal features described previously in literature^3^ and by the cell line vendor (DSMZ) (**Figure 1**). G-banding confirmed the majority of the large chromosomal aberrations expected in the REH cell line, including the deletion of one X chromosome, a deletion on the short arm of chromosome 3, and the balanced t(5;12). Our G-banding, however, did not completely resolve the four-way t(4;12;21;16). A comparison of the analog karyotypes is available in the Supplemental Discussion and Supplemental Table S2.

**Figure 1.**
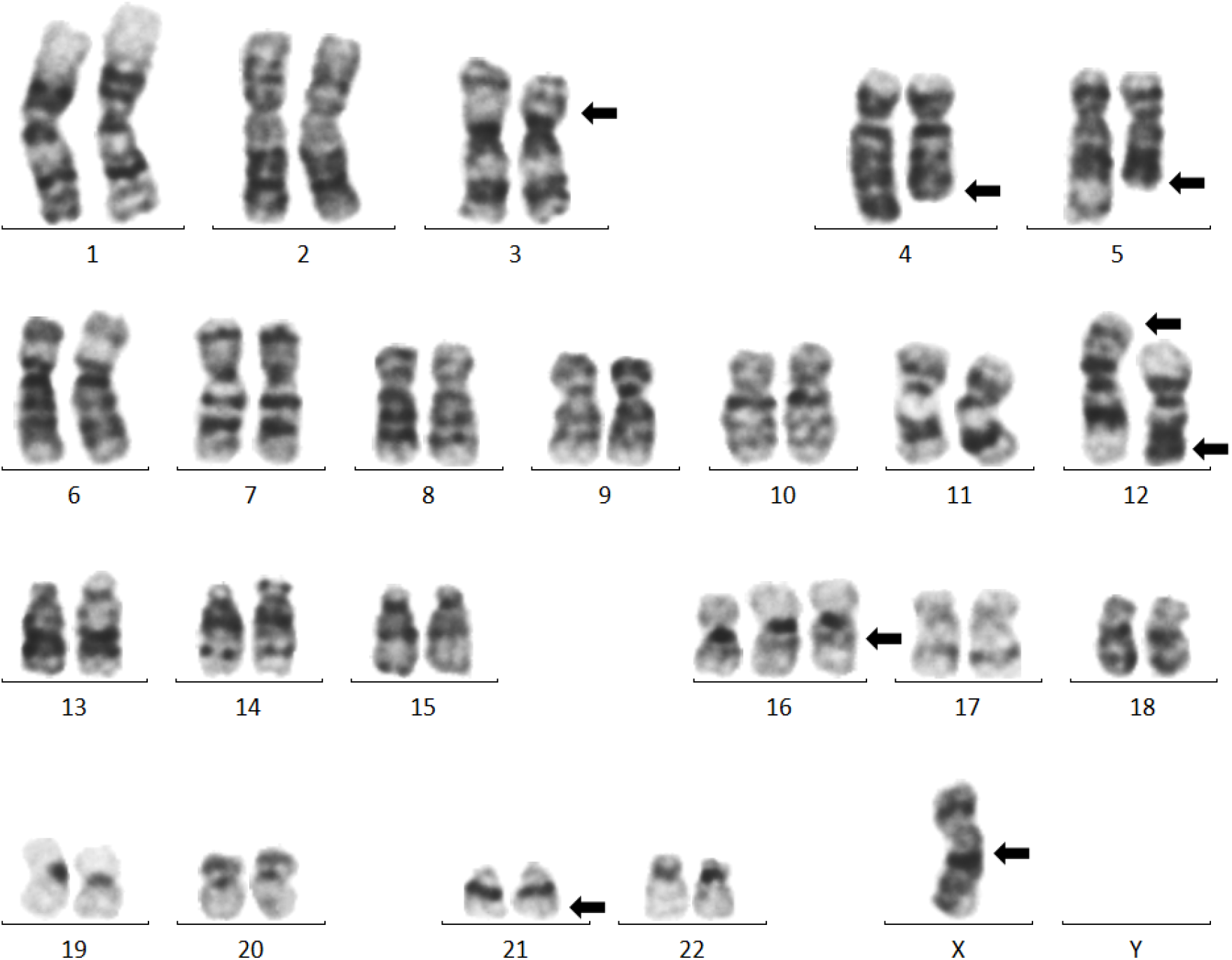
G-banding of the REH cell line, used to verify the karyotype provided by the cell line vendor. Arrows mark the chromosomes with visible aberrations, reflecting the major features of the stemline described in the DSMZ karyotype: *46(44-47)<2n>X, -X, +16, del(3)(p22), t(4;12;21;16)(q32;p13;q22;q24.3)-inv(12)(p13q22), t(5;12)(q31-q32;p12), der(16)t(16;21)(q24.3;q22) - sideline with inv(5)der(5)(p15q31),+18*. G-banding showed that the cells in the present study did not contain the sideline.

### The SV landscape stratified by size

We generated three SV callsets by running Sniffles on PacBio and ONT long-read WGS data, and TIDDIT on Illumina short-read WGS data (**Table 1**). In total, TIDDIT generated 5726 unfiltered SV candidates from the Illumina data, while Sniffles generated 36121 candidates from PacBio and 36648 candidates from ONT. Of SVs with length > 100bp, there were 10262 variants in the long-read consensus set (called in both ONT and PacBio), while the three-way consensus set (intersection of all three WGS callsets) contained 2072 variants. The non-translocation SVs were binned by size into Small (100 bp −1 kbp), Medium (1 kbp - 10 kbp) and Large (> 10 kbp); in these categories, the long-read consensus sets contained 8993, 1420, and 19 variants, respectively, while the three-way consensus sets contained 1485, 414 and 34 variants. (**Figure 2**).

**Figure 2.**
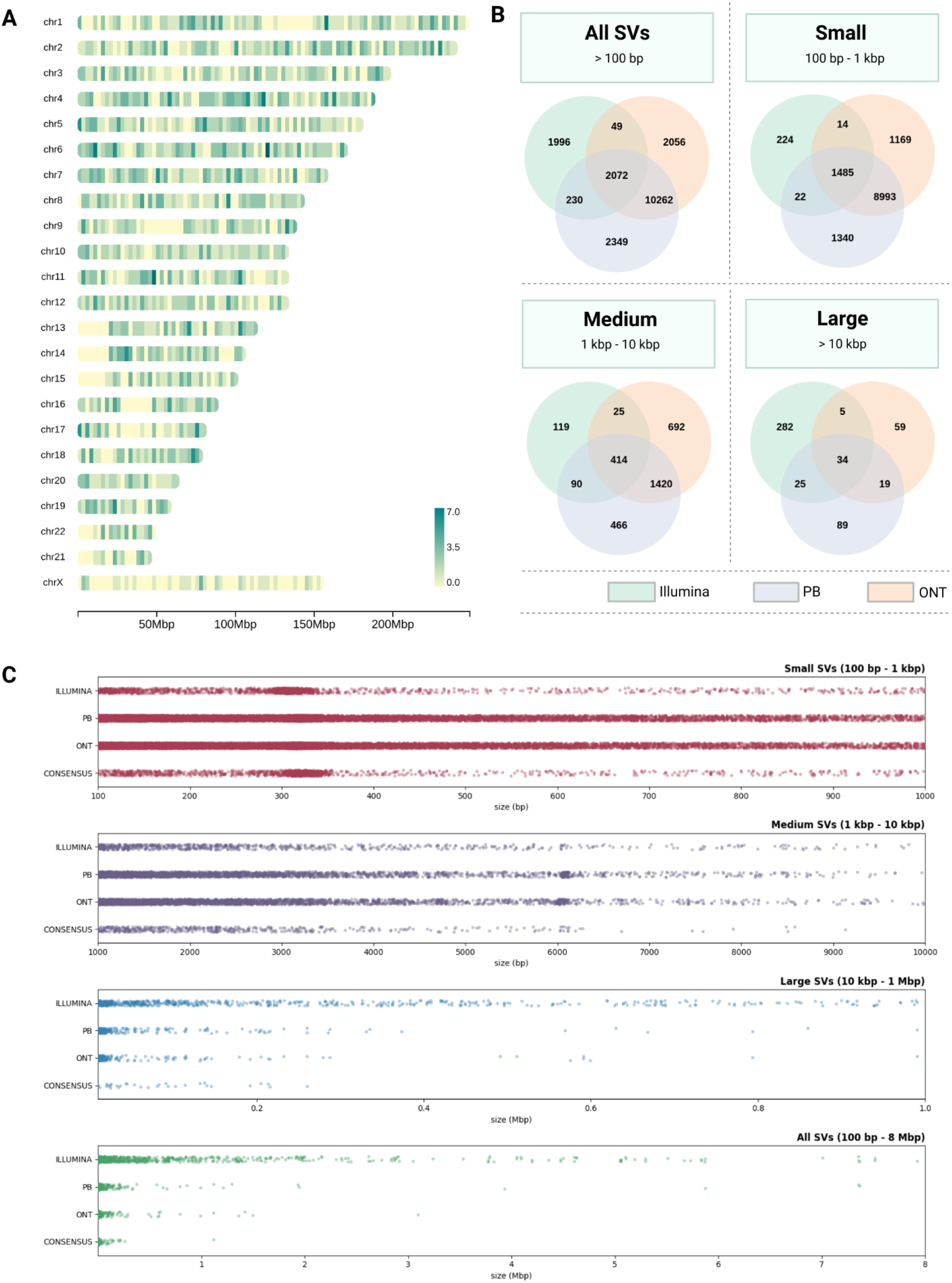
REH structural variants detected across PacBio, ONT and Illumina SV callsets. a) chromosomal heatmap of the three-way consensus callset, showing the total number of SV calls at each locus b) Venn diagram showing the the number of SV calls found to overlap in each combination of callsets c) strip plots showing SVs from each callset, stratified by size.

After automated filtering for large-scale SV’s (> 100 kbp) and interchromosomal translocations (see Supplemental Methods), there remained: 694 SV candidates from Illumina; 127 from PacBio; and 35 from ONT. All remaining Sniffles candidates were manually inspected in IGV, while a pseudo-random sampling from the TIDDIT set selected 29 candidates across 16 chromosomes for further inspection.

### Refinement of known REH aberrations at base-pair resolution

We detected the established and expected chromosomal aberrations in each of the three SV callsets (**Table 2**). These features include a deleted chromosome X, gain of chromosome 16; a 26mbp del(3)(p22.3;p14.2); a balanced translocation between chromosomes 5 and 12; and finally, a complex four-way rearrangement between chromosomes 4, 12, 21, and 16, resulting in the *ETV6-RUNX1* fusion gene. We were also able to determine that the t(5;12) and the four-way translocation involve two different alleles of chromosome 12, as the two different p-arms of chromosome 12 (from p13.2 to the telomere) were found to be fused to either chromosome 5 or 21. Combining the different SV callsets facilitated the resolution of these breakpoints down to a base-pair accuracy, thereby resolving inconsistencies in the previously documented REH karyotypes (see Supplemental Discussion and Supplemental Table S2).

**Table 2.**
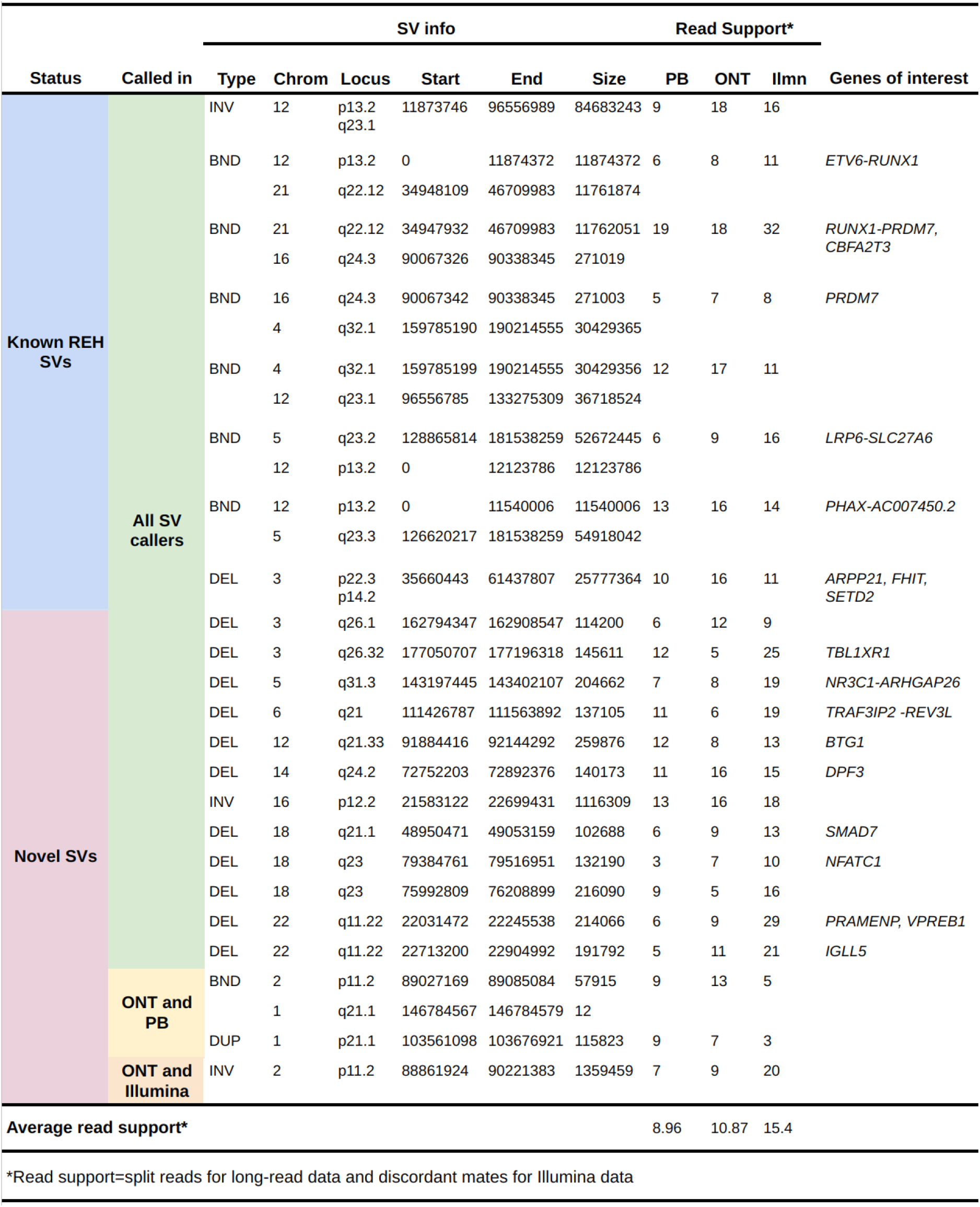
The structural variants observed in REH across the different SV callsets.

We reconstructed the molecular events of the four-way translocation as follows. The p-arm of chromosome 12 (12Mbp), with breakpoint at p13.2 in intron 5 of *ETV6*, was translocated to chr21q22.12, resulting in the canonical, subtype-defining fusion gene *ETV6-RUNX1*. The q-arm of chromosome 21 (12Mbp), with breakpoint at q22.12 in intron 1 of *RUNX1*, was translocated to chr16q24.3. The q-arm of chromosome 16 (271kbp), with a breakpoint at q24.3 in intron 4 of *PRDM7*, was translocated to chr4q32.1. Finally, the q-arm of chromosome 4 (30Mbp), with breakpoint at q32.1, was translocated to chr12q23.1 (where chromosome 12 has undergone an 85Mbp inversion between p13.2 and q23.1), completing the circular exchange.

We were able to detect known aneuploidies: a deletion of chromosome X was detected in all data sets by a marked decrease in the average depth of coverage (DC), while an additional chromosome 16 was detected by a corresponding increase (Supplemental Table S3). All three analog karyotypes indicate two copies of the derived chromosome 16 resulting from the translocation t(16;21)(q24;q22); we confirmed this through DC and analysis of reads mapping to both chromosomes 16 and 21 (see Supplemental Discussion). While this evidence supports the presence of two copies of der(16)t(16;21), we found a single set of breakends for this translocation, suggesting that this aneuploidy was caused by an aberrant mitotic event occurring after the translocation.

### Large-scale SVs discovered in the REH genome

In total, we identified 16 intrachromosomal and seven interchromosomal SVs (**Figure 3**), of which one interchromosomal translocation and 14 intrachromosomal SVs (deletions, duplications and inversions > 100 kbp) were previously unidentified in the analog karyotypes (**Table 2**). We identified a pair of rearrangements on chromosome 2 including a 1.4 Mbp inversion at p11.2 and an adjacent, unbalanced 58 kb translocation resulting in derived chromosome der(1)t(1;2)(q21.1;p11.2). Our findings also include a 116 kbp duplication on chromosome 1p21.1, a 1.1 Mbp inversion on chromosome 16p12.2, and deletions on seven different chromosomes. These include two deletions on chromosome 3 at q26.1 (114kbp) and q26.32 (146bkp), the latter deleting exons 2-5 of gene *TBL1XR1*, which has been implicated in GC drug resistance ^44^. A deletion on chromosome 5q31.3 (205kbp) was found to focally delete exons 2-9 of GC receptor gene *NR3C1* ^45^, while a deletion on chromosome 12q21.33 (260kbp) was found in the region encoding the ALL-associated gene *BTG1* ^46^. Three deletions were found on chromosome 18, one at q21.1 and two at q23, including the 132kbp deletion affecting exon 10 of the leukemia-associated gene *NFATC1* ^47^. Finally, two deletions were discovered on chromosome 22q11.22, of which a 214 kbp variant was found to entirely delete the gene *VPREB1*, which encodes the surrogate light chain involved in the formation of pre-B cell receptor (pre-BCR) and whose genetic loss contributes to leukemogenesis ^48^ (**Supplemental Figure 1-5**).

**Figure 3.**
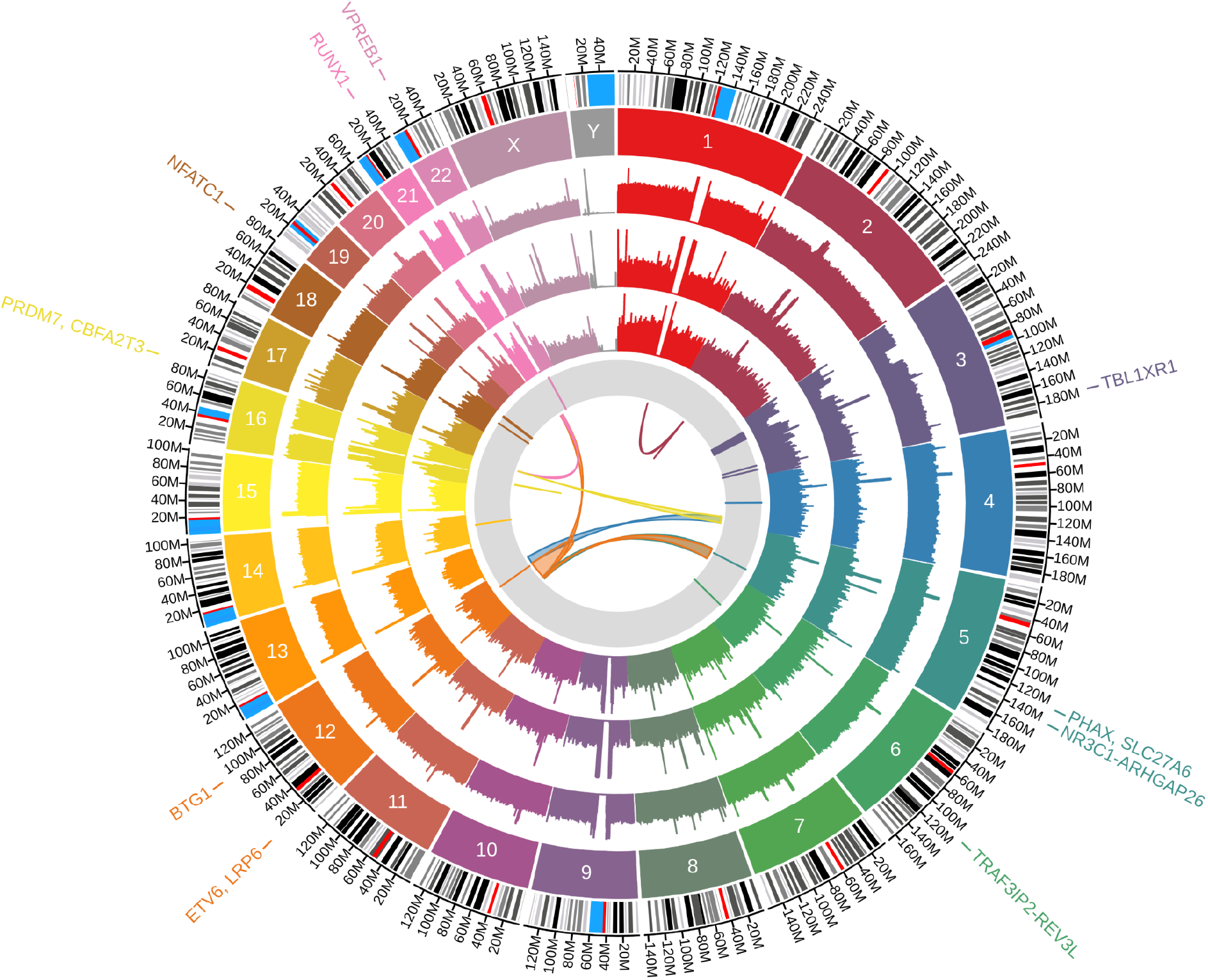
Large-scale REH structural variants detected across PacBio, ONT and Illumina SV callsets. The innermost circle shows inversions and interchromosomal translocations. The gray band depicts deletions > 100 kb. The rainbow bands show the depth of coverage in PacBio, ONT, and Illumina data sets, respectively, followed by chromosome number. The outermost band indicates the GRCh38 reference cytoband with genes disrupted listed on the outer edge of the plot.

Of the 23 confirmed SVs, a total of 20 were called in all three SV callsets, including all aberrations known from the analog karyotypes and 12 of the novel SVs. An additional two SVs, in repetitive regions of chromosomes 1 and 2, were called in both PacBio and ONT, but not in Illumina data, while one SV was called in both Illumina and ONT. All of the confirmed SVs were supported by at least three split reads from both ONT and PacBio, and at least three discordant read pairs from Illumina, with an average read support of 9.0 from PacBio, 10.9 from ONT and 15.4 from Illumina. (**Table 2**). Additional details can be found in the Supplemental Discussion and Table S4.

### Seven expressed fusion genes, including two in-frame fusions

We used short- and long-read RNA-sequencing data together with seven fusion gene calling softwares to detect potential fusion genes in REH. In total, the unfiltered fusion callset consisted of 11099 fusion gene candidates. After stringent filtering (see Supplemental Methods), 30 candidates remained. Additional manual examination of split and spanning reads, as well as discordant mate pairs, retained seven high-confidence fusion candidates whose breakpoints could be confirmed in IGV with supporting genomic breakpoints in the WGS data (**Figure 4**).

**Figure 4.**
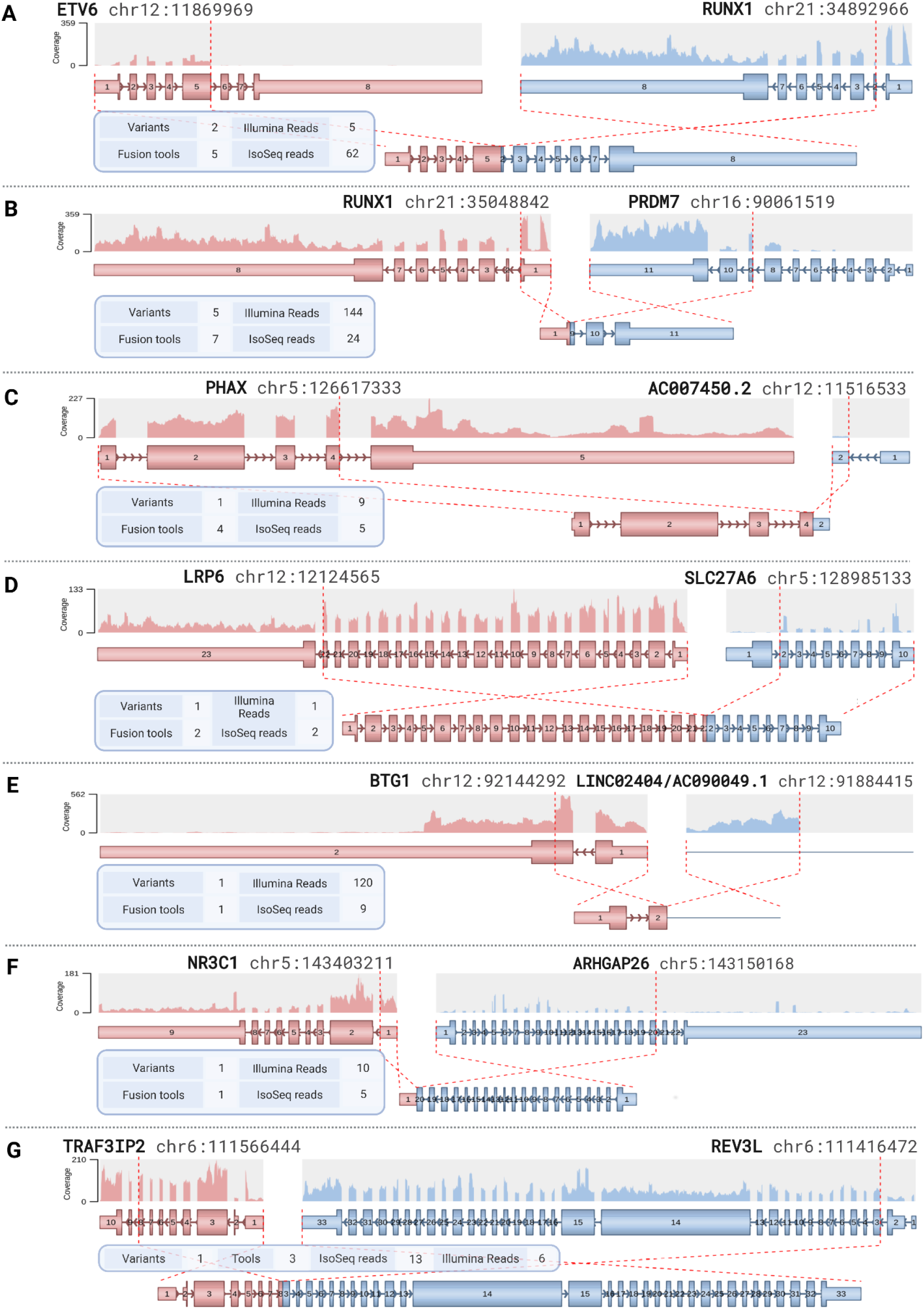
REH fusion gene breakpoints, visualized by the Arriba module of the nf-core/rnafusion pipeline. a) the more highly expressed of the two splice variants of *ETV6-RUNX1*, resulting from t(12;21) b) the most highly expressed of the five splicing variants of *RUNX1-PRDM7*, from t(16;21) c) *PHAX-AC007450.2* and d) *LRP6-SLC27A6*, both from t(5;12) e) *BTG1-LINC02404/AC090049.17*, from del(12)(q21.33), f) *NR3C1-ARHGAP26*, from del(5)(q31.3) g) *TRAF3IP2-REV3L*, from del(6)(q21)

Out of the seven confirmed fusion genes, two were in-frame. First, the expected *ETV6-RUNX1* fusion, resulting from the canonical subtype-defining translocation t(12;21), was detected as two splicing variants. Transcripts of both variants retained the Sterile alpha motif (SAM)/Pointed domain from *ETV6*, and the Runt and Runx inhibition domains from *RUNX1* (**Figure 4a**). Five splicing variants of a second in-frame fusion gene, *RUNX1-PRDM7*, resulted from the t(16;21) occurring in two copies of the der(16). In the splicing variant involving exon 5 of *PRDM7*, the *RUNX1-PRDM7* fusion retained the promoter from *RUNX1* and the SSXRD motif from *PRDM7* (**Figure 4b**). Two out-of-frame fusion genes were found to arise from the balanced t(5;12): *PHAX-AC007450.2* and *LRP6-SLC27A6* (**Figure 4c, 4d**). A fusion transcript arose from the 260kbp deletion del(12)(q21.33), which resulted in a truncated *BTG1* gene, with a breakpoint in exon 2. In this fusion, the truncated *BTG1* transcript was fused with an expressed non-coding region originating between two long non-coding RNA genes *LINC02404* and *AC090049.1*. (**Figure 4e**). Finally, the 205 kbp del(5)(q31.3) resulted in the antisense transcript *NR3C1-ARHGAP26* (**Figure 4f**), while the 137kb del(6)q21 resulted in the fusion gene *TRAF3IP2-REV3L* (**Figure 4g**). Additional details can be found in the Supplemental Discussion and Supplemental Table S5.

Of note, three of the seven confirmed fusion genes involve the partner genes *ETV6* and *RUNX1* or regions in their immediate vicinity (*ETV6-RUNX1, PHAX-AC007450.2 and RUNX1-PRDM7*). We found that only one allele of chromosome 21 was involved in both *RUNX1* fusions, while both alleles of *ETV6* were involved in different chromosomal aberrations occurring on chromosome 12, with one allele forming the *ETV6-RUNX1* fusion and the other allele undergoing a 584kb deletion between the breaking and fusion events of the t(5;12), deleting the entire gene and leaving no wild-type *ETV6* in the genome.

The performance of the seven fusion gene calling tools varied widely, finding from 31 to 5520 fusion gene candidates, with a sensitivity ranging from 14.29% to 100%, and a false positive rate (FPR) of 50% to 99.94%. Additional details can be found in the Supplemental Discussion and Table S6.

### A comprehensive karyotype

Our integrative short- and long-read genomic and transcriptomic analysis allows us to provide a revised, comprehensive digital karyotype for the REH cell line: *46, X, -X, dup(1)(p21.1), t(1;2)(q21.1;p11.2)-inv(2)(p11.2), del(3)(p14.2p22.3), del(3)(q26.1), del(3)(q26.32), t(4;12;21;16)(q32.1;p13.2;q22.12;q24.3)-inv(12)(p13.2q23.1), del(5)(q31.3), t(5;12)(q23.2-q23.3;p13.2), del(6)(q21), del(12)(q21.33), del(14)(q24.2), inv(16)(p12.2), der(16)t(16;21)(q24.3;q22.12) x 2, del(18)(q21.1), del(18)(q23) x 2, del(22)(q11.22) x 2* (**Figure 5).**

**Figure 5.**
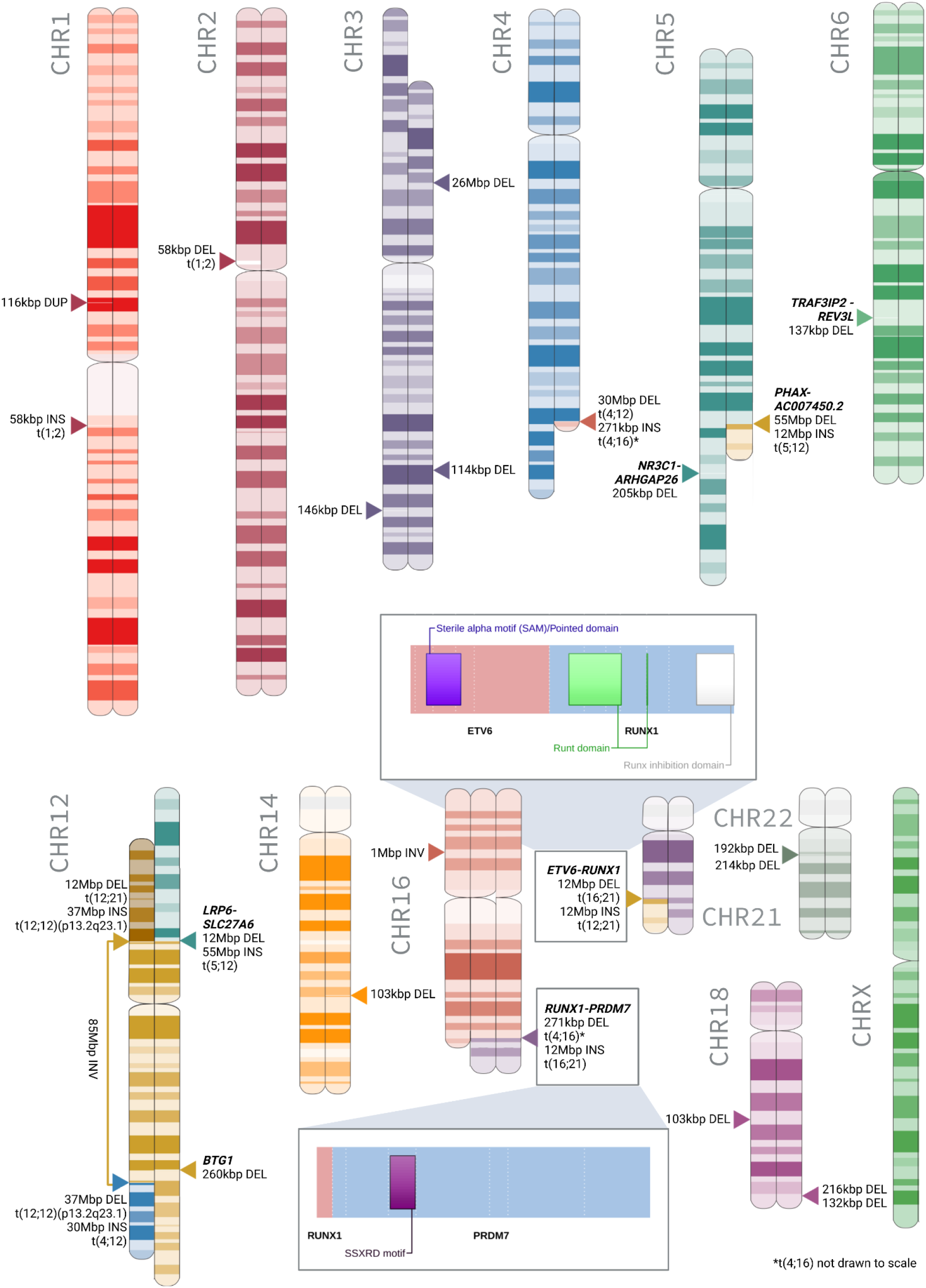
The aberrant chromosomes of the comprehensive REH karyotype. Highlighted are the retained protein domains for *ETV6-RUNX1* and *RUNX1-PRDM7*, the two REH in-frame fusion genes.

## DISCUSSION

Cell lines continue to be a vital resource for functional studies. Several well-established cancer cell lines are highly rearranged, with underreported genomic complexity ^49–51^. Despite its frequent use for functional and mechanistic studies in leukemia research ^8–10,52^, a comprehensive analysis of the complex REH genome has never been undertaken. Herein, we used a combination of short- and long-read DNA and RNA sequencing to provide a complete digital karyotype for the REH cell line. This clarifies the ambiguity left by traditional karyotyping and resolves a number of known and novel aberrations in the REH genome, which are of potential biological importance.

The REH cell line comes from a relapsed patient, making it particularly notable that several deletions identified in the present study, affecting genes *TBL1XR1* (146k deletion on 3q26.32), *NR3C1* (205 kbp deletion on 5q31.3) and *BTG1* (260kbp deletion on 12q21.33), have been shown to contribute to its GC drug resistance ^44–46^. GCs are a key component of ALL treatment protocols and the roles of *NR3C1* and *BTG1* expression and perturbation have been widely analyzed in the REH cell line ^46,53,54^. Despite the many functional studies that use REH, few have addressed the genomic aberrations underlying its phenotype. The *BTG1* deletion identified in the present study has been previously noted in analyses of REH using arrays ^55^ and PCR ^46^, while *NR3C1* has an established nonsense (p.Gln528Ter) mutation on one allele ^56^ and a deletion on the other ^57^. Recurrent deletions of *TBL1XR1*, which encodes for nuclear hormone receptor co-repressors, have been shown to contribute to dysregulated gene expression and GC resistance in *ETV6-RUNX1* positive ALL, including REH^58,59^.

In the present study we identified a previously unknown REH variant (214 kbp deletion on 22q11.22) that deletes the *VPREB1* gene, which codes for part of the surrogate light chain that forms pre-BCRs. *VPREB1* deletions have previously been shown to occur in up to 40% of *ETV6-RUNX1* positive ALL patients at diagnosis and >70% at relapse^60^, contributing to the disordered light chain rearrangement that acts as a leukemogenic trigger ^48^. The *VPREB1* deletion in REH, which also encompasses part of the pseudogene *PRAMENP*, is adjacent to another 192 kbp deletion, affecting the chronic lymphocytic leukemia-associated gene *IGLL5^61^*; the gene that lies between these two deletions, *PRAME*, has been identified as a potential therapeutic target for pediatric ALL^62^. Recurrent deletions at the 22q11.22 locus have been previously identified as a prognostic marker of poor outcome for certain B-ALL patients^63^, making this a genomic region warranting further research.

Likewise, the long arm of chromosome 18 was the site of recurring deletions. Of the three large-scale deletions found between 18q21-q23, one was found to affect the *NFATC1* gene (132kb del on 18q23), a member of the *NFAT* family involved in the calcineurin signaling pathway, which has been identified as a potential therapeutic target ^47^. In addition, the 26 Mbp deletion on chromosome 3 encompasses a large number of genes, including *SETD2*, which is associated with chemotherapy resistance^64^*; ARPP21*, a frequent deletion in *ETV6-RUNX1* -like ALL cases^65^, and *FHIT*, a tumor suppressor gene that has been shown to be abnormally expressed in ALL^66^.

The fusion gene arising from the unbalanced t(16;21)(q24;q22) is *RUNX1-PRDM7*. This highly expressed fusion occurs on the same *RUNX1* allele as the canonical *ETV6-RUNX1* fusion. In contrast to previous reports ^67,68^, our data indicate that this fusion is in-frame, retaining the SSXRD motif of *PRDM7*. Because of the close sequence similarity between *PRDM7* (chromosome 16) and its paralogous copy *PRDM9* (chromosome 5)^69^, this fusion was misidentified as *RUNX1-PRDM9* by two short-read fusion callers and a mismapping of up to 127 short RNA-seq reads. *PRDM9*, which shares the SSXRD, SET and KRAB protein domains with *PRDM7*, has been implicated in genetic predisposition to pediatric B-ALL ^70,71^, while our findings suggest that this association might extend to other members of the *PRDM* gene family.

In assessing the performance of the different long- and short-read technologies, we found that it was beneficial to implement an integrative process using a variety of tools and datasets due to their distinct strengths in detecting variants of different types and sizes ^72^. Overall, we found that the ONT dataset was the most useful for SV discovery, with the highest sensitivity and lowest FPR; however, when adjusting for the slightly lower DC, PacBio had a comparable performance. Importantly, both long-read datasets were able to resolve variants in highly repetitive regions that were missed in the short-read data.

We found that SV detection using TIDDIT on Illumina short reads resulted in an FPR of over 98%, limiting its standalone utility in SV discovery in the absence of a matched normal control; in particular, a known problem with this method is a high rate of falsely identified translocations ^32^. However, short, paired reads were useful in confirming and resolving breakpoints suggested by the long-reads; thus, given the high throughput and low cost, short reads remain a sensible choice for clinical screenings targeting specific variants. Of note, the high FPR and low sensitivity observed in this study are the statistical side effects of using a consensus method combining SV calls from several different methods; the upside to this approach is high precision ^24^.

In assessing fusion detection capabilities, we found short-read RNA-seq data to be more informative than the IsoSeq dataset, largely due to the superior performance of the short-read fusion caller Arriba, which has been shown to outperform both other short-read fusion callers^73^, and long-read callers in low-coverage contexts^74^.

In summary, our findings highlight the need for comprehensive genomic analysis of commonly used and perturbed cell lines. Further, we underscored the importance of the increased informational context of long reads and how they can identify critical biologically important SVs. Long reads hold promise to more accurately capture the complexity of SVs in cancer genomes and these results add to the growing evidence for the potential value of long-read sequencing in clinical cancer genomics. Finally, we have cataloged multiple aberrations in REH that are frequently observed in high-risk ALL patients, making this cell line a highly relevant model for researchers investigating the biological underpinnings of relapse and GC resistance.

## ACKNOWLEDGEMENTS

This project was funded by the Swedish Research Council (#2019-01976), the Swedish Childhood Cancer Fund (#2019-0046), and the Göran Gustafsson Foundation. Sequencing was performed at the National Genomics Infrastructure (NGI) at SciLifelab in Uppsala. NGI is funded by SciLifeLab, the Swedish Research Council RFI, and the Knut and Alice Wallenberg Foundation. Computational resources were provided by the Swedish National Infrastructure for Computing (SNIC), also funded by the Swedish Research Council. The authors would like to acknowledge Elin Övernäs and Johanna Lagensjö for their assistance with short-read sequencing, Sara Ekberg, Pontus Larsson, Rikard Erlandsson and Susanne Reinsbach for bioinformatics support and Mai-Britt Mosbech, Susana Häggkvist and Anna Petri for assistance with high molecular weight DNA extraction and long-read sequencing.

## AUTHORSHIP CONTRIBUTIONS

MLW, AA, LF and JN conceived the research and constructed the experimental design. JN and LF acquired funding. MLW, GA, IB, YMZ, AL, and AA analyzed data. HG and AR provided material and sequencing libraries. AB and YMZ provided expertise on karyotyping. MLW made the figures. MLW and JN wrote the manuscript. All authors have read and agreed to the published version of the manuscript.

## REFERENCES

1. Matsuo, Y. & Drexler, H. G. Establishment and characterization of human B cell precursor-leukemia cell lines. Leuk. Res. 22, 567–579 (1998).

2. Rosenfeld, C. et al. Phenotypic characterisation of a unique non-T, non-B acute lymphoblastic leukaemia cell line. Nature 267, 841–843 (1977).

3. Uphoff, C. C. et al. Occurrence of TEL-AML1 fusion resulting from (12;21) translocation in human early B-lineage leukemia cell lines. Leukemia 11, 441–447 (1997).

4. Bachmann, P. S. et al. Divergent Mechanisms of Glucocorticoid Resistance in Experimental Models of Pediatric Acute Lymphoblastic Leukemia. Cancer Res. 67, 4482–4490 (2007).

5. Wall, N. R., Mohammad, R. M. & Al-Katib, A. M. Bax:Bcl-2 ratio modulation by bryostatin 1 and novel antitubulin agents is important for susceptibility to drug induced apoptosis in the human early pre-B acute lymphoblastic leukemia cell line, Reh. Leuk. Res. 23, 881–888 (1999).

6. Jiang, N. et al. Identification of prognostic protein biomarkers in childhood acute lymphoblastic leukemia (ALL). J. Proteomics 74, 843–857 (2011).

7. Tzoneva, G. et al. Clonal evolution mechanisms in NT5C2 mutant-relapsed acute lymphoblastic leukaemia. Nature 553, 511–514 (2018).

8. Cousins, A. et al. Central nervous system involvement in childhood acute lymphoblastic leukemia is linked to upregulation of cholesterol biosynthetic pathways. Leukemia 36, 2903–2907 (2022).

9. Leo, I. R. et al. Integrative multi-omics and drug response profiling of childhood acute lymphoblastic leukemia cell lines. Nat. Commun. 13, 1691 (2022).

10. Wray, J. P. et al. Regulome analysis in B-acute lymphoblastic leukemia exposes Core Binding Factor addiction as a therapeutic vulnerability. Nat. Commun. 13, 7124 (2022).

11. Jobanputra, V. et al. Application of ROMA (representational oligonucleotide microarray analysis) to patients with cytogenetic rearrangements. Genet. Med. 7, 111–118 (2005).

12. Dwivedi, A. C. nee et al. Clinical utility of chromosomal microarray analysis in the diagnosis and management of monosomy 7 mosaicism. Mol. Cytogenet. 7, 93 (2014).

13. Tsang, E. S. et al. Uncovering Clinically Relevant Gene Fusions with Integrated Genomic and Transcriptomic Profiling of Metastatic Cancers. Clin. Cancer Res. 27, 522–531 (2021).

14. Berglund, E. et al. A Study Protocol for Validation and Implementation of Whole-Genome and -Transcriptome Sequencing as a Comprehensive Precision Diagnostic Test in Acute Leukemias. Front. Med. 9, 842507 (2022).

15. Cuppen, E. et al. Implementation of Whole-Genome and Transcriptome Sequencing Into Clinical Cancer Care. JCO Precis. Oncol. e2200245 (2022) doi:10.1200/PO.22.00245.

16. Amarasinghe, S. L. et al. Opportunities and challenges in long-read sequencing data analysis. Genome Biol. 21, 30 (2020).

17. Xiao, W. et al. Toward best practice in cancer mutation detection with whole-genome and whole-exome sequencing. Nat. Biotechnol. 39, 1141–1150 (2021).

18. Jobanputra, V. et al. Clinical interpretation of whole-genome and whole-transcriptome sequencing for precision oncology. Semin. Cancer Biol. 84, 23–31 (2022).

19. Roberts, H. E. et al. Short and long-read genome sequencing methodologies for somatic variant detection; genomic analysis of a patient with diffuse large B-cell lymphoma. Sci. Rep. 11, 6408 (2021).

20. Meggendorfer, M. et al. Analytical demands to use whole-genome sequencing in precision oncology. Semin. Cancer Biol. 84, 16–22 (2022).

21. Ebbert, M. T. W. et al. Systematic analysis of dark and camouflaged genes reveals disease-relevant genes hiding in plain sight. Genome Biol. 20, 97 (2019).

22. Ardui, S., Ameur, A., Vermeesch, J. R. & Hestand, M. S. Single molecule real-time (SMRT) sequencing comes of age: applications and utilities for medical diagnostics. Nucleic Acids Res. 46, 2159–2168 (2018).

23. Hu, T., Chitnis, N., Monos, D. & Dinh, A. Next-generation sequencing technologies: An overview. Hum. Immunol. 82, 801–811 (2021).

24. Liu, Y. et al. Comparison of multiple algorithms to reliably detect structural variants in pears. BMC Genomics 21, 61 (2020).

25. Conlin, L. K., Aref-Eshghi, E., McEldrew, D. A., Luo, M. & Rajagopalan, R. Long-read sequencing for molecular diagnostics in constitutional genetic disorders. Hum. Mutat. 43, 1531–1544 (2022).

26. Di Tommaso, P. et al. Nextflow enables reproducible computational workflows. Nat. Biotechnol. 35, 316–319 (2017).

27. Ewels, P. A. et al. The nf-core framework for community-curated bioinformatics pipelines. Nat. Biotechnol. 38, 276–278 (2020).

28. Garcia, M. et al. Sarek: A portable workflow for whole-genome sequencing analysis of germline and somatic variants. F1000Research 9, 63 (2020).

29. Eisfeldt, J., Vezzi, F., Olason, P., Nilsson, D. & Lindstrand, A. TIDDIT, an efficient and comprehensive structural variant caller for massive parallel sequencing data. F1000Research 6, 664 (2017).

30. Li, H. Minimap2: pairwise alignment for nucleotide sequences. Bioinformatics 34, 3094–3100 (2018).

31. Danecek, P. et al. Twelve years of SAMtools and BCFtools. GigaScience 10, giab008 (2021).

32. Sedlazeck, F. J. et al. Accurate detection of complex structural variations using single-molecule sequencing. Nat. Methods 15, 461–468 (2018).

33. Jeffares, D. C. et al. Transient structural variations have strong effects on quantitative traits and reproductive isolation in fission yeast. Nat. Commun. 8, 14061 (2017).

34. Amemiya, H. M., Kundaje, A. & Boyle, A. P. The ENCODE Blacklist: Identification of Problematic Regions of the Genome. Sci. Rep. 9, 9354 (2019).

35. Robinson, J. T., Thorvaldsdóttir, H., Wenger, A. M., Zehir, A. & Mesirov, J. P. Variant Review with the Integrative Genomics Viewer. Cancer Res. 77, e31–e34 (2017).

36. Proks, M. et al. nf-core/rnafusion: nf-core/rnafusion:1.2.0. (2020) doi:10.5281/ZENODO.3946477.

37. Uhrig, S. et al. Accurate and efficient detection of gene fusions from RNA sequencing data. Genome Res. 31, 448–460 (2021).

38. Nicorici, D. et al. FusionCatcher-a tool for finding somatic fusion genes in paired-end RNA-sequencing data. http://biorxiv.org/lookup/doi/10.1101/011650 (2014) doi:10.1101/011650.

39. Melsted, P. et al. Fusion detection and quantification by pseudoalignment. http://biorxiv.org/lookup/doi/10.1101/166322 (2017) doi:10.1101/166322.

40. Ma, C., Shao, M. & Kingsford, C. SQUID: transcriptomic structural variation detection from RNA-seq. Genome Biol. 19, 52 (2018).

41. Haas, B. J. et al. Accuracy assessment of fusion transcript detection via read-mapping and de novo fusion transcript assembly-based methods. Genome Biol. 20, 213 (2019).

42. Tardaguila, M. et al. SQANTI: extensive characterization of long-read transcript sequences for quality control in full-length transcriptome identification and quantification. Genome Res. 28, 396–411 (2018).

43. Davidson, N. M., Majewski, I. J. & Oshlack, A. JAFFA: High sensitivity transcriptome-focused fusion gene detection. Genome Med. 7, 43 (2015).

44. Jones, C. L. et al. Loss of TBL1XR1 Disrupts Glucocorticoid Receptor Recruitment to Chromatin and Results in Glucocorticoid Resistance in a B-Lymphoblastic Leukemia Model. J. Biol. Chem. 289, 20502–20515 (2014).

45. Xiao, H. et al. Haploinsufficiency of NR3C1 drives glucocorticoid resistance in adult acute lymphoblastic leukemia cells by down-regulating the mitochondrial apoptosis axis, and is sensitive to Bcl-2 blockage. Cancer Cell Int. 19, 218 (2019).

46. van Galen, J. C. et al. BTG1 regulates glucocorticoid receptor autoinduction in acute lymphoblastic leukemia. Blood 115, 4810–4819 (2010).

47. Medyouf, H. & Ghysdael, J. The calcineurin/NFAT signaling pathway: A NOVEL therapeutic target in leukemia and solid tumors. Cell Cycle 7, 297–303 (2008).

48. Mangum, D. S. et al. VPREB1 deletions occur independent of lambda light chain rearrangement in childhood acute lymphoblastic leukemia. Leukemia 28, 216–220 (2014).

49. Nattestad, M. et al. Complex rearrangements and oncogene amplifications revealed by long-read DNA and RNA sequencing of a breast cancer cell line. Genome Res. 28, 1126–1135 (2018).

50. Arora, K. et al. Deep whole-genome sequencing of 3 cancer cell lines on 2 sequencing platforms. Sci. Rep. 9, 19123 (2019).

51. Aganezov, S. et al. Comprehensive analysis of structural variants in breast cancer genomes using single-molecule sequencing. Genome Res. 30, 1258–1273 (2020).

52. Diedrich, J. D. et al. Profiling chromatin accessibility in pediatric acute lymphoblastic leukemia identifies subtype-specific chromatin landscapes and gene regulatory networks. Leukemia 35, 3078–3091 (2021).

53. Liu, H. et al. Association Between NR3C1 Mutations and Glucocorticoid Resistance in Children With Acute Lymphoblastic Leukemia. Front. Pharmacol. 12, 634956 (2021).

54. van der Zwet, J. C. G., Smits, W., Buijs-Gladdines, J. G. C. A. M., Pieters, R. & Meijerink, J. P. P. Recurrent NR3C1 Aberrations at First Diagnosis Relate to Steroid Resistance in Pediatric T-Cell Acute Lymphoblastic Leukemia Patients. HemaSphere 5, e513 (2021).

55. Tsuzuki, S. et al. Genetic abnormalities involved in t(12;21) Tel–AML1 acute lymphoblastic leukemia: Analysis by means of array-based comparative genomic hybridization. Cancer Sci. 98, 698–706 (2007).

56. Tamai, M. et al. Glucocorticoid receptor gene mutations confer glucocorticoid resistance in B-cell precursor acute lymphoblastic leukemia. J. Steroid Biochem. Mol. Biol. 218, 106068 (2022).

57. Grausenburger, R. et al. Genetic alterations in glucocorticoid signaling pathway components are associated with adverse prognosis in children with relapsed *ETV6/RUNX1* -positive acute lymphoblastic leukemia. Leuk. Lymphoma 57, 1163–1173 (2016).

58. Parker, H. et al. The complex genomic profile of *ETV6-RUNX1* positive acute lymphoblastic leukemia highlights a recurrent deletion of TBL1XR1. Genes. Chromosomes Cancer 47, 1118–1125 (2008).

59. Jones, C. L. et al. Deletions In TBL1XR1 Results In Glucocorticoid Resistance By Decreasing Glucocorticoid Signaling In Childhood B-Lymphoblastic Leukemia. Blood 122, 602–602 (2013).

60. Kuster, L. et al. ETV6/RUNX1-positive relapses evolve from an ancestral clone and frequently acquire deletions of genes implicated in glucocorticoid signaling. Blood 117, 2658–2667 (2011).

61. Kasar, S. et al. Whole-genome sequencing reveals activation-induced cytidine deaminase signatures during indolent chronic lymphocytic leukaemia evolution. Nat. Commun. 6, 8866 (2015).

62. Steinbach, D., Viehmann, S., Zintl, F. & Gruhn, B. PRAME gene expression in childhood acute lymphoblastic leukemia. Cancer Genet. Cytogenet. 138, 89–91 (2002).

63. Mangum, D. S. et al. Association of Combined Focal 22q11.22 Deletion and IKZF1 Alterations With Outcomes in Childhood Acute Lymphoblastic Leukemia. JAMA Oncol. 7, 1521 (2021).

64. Mar, B. G. et al. SETD2 alterations impair DNA damage recognition and lead to resistance to chemotherapy in leukemia. Blood 130, 2631–2641 (2017).

65. Zaliova, M. et al. ETV6/RUNX1 -like acute lymphoblastic leukemia: A novel B-cell precursor leukemia subtype associated with the CD27/CD44 immunophenotype: ZALIOVA et al. Genes. Chromosomes Cancer 56, 608–616 (2017).

66. Hallas, C. et al. Loss of FHIT Expression in Acute Lymphoblastic Leukemia1. Clin. Cancer Res. 5, 2409–2414 (1999).

67. Lilljebjörn, H. et al. RNA-seq identifies clinically relevant fusion genes in leukemia including a novel MEF2D/CSF1R fusion responsive to imatinib. Leukemia 28, 977–979 (2014).

68. Ghandi, M. et al. Next-generation characterization of the Cancer Cell Line Encyclopedia. Nature 569, 503–508 (2019).

69. Fumasoni, I. et al. Family expansion and gene rearrangements contributed to the functional specialization of PRDM genes in vertebrates. BMC Evol. Biol. 7, 187 (2007).

70. Hussin, J. et al. Rare allelic forms of *PRDM9* associated with childhood leukemogenesis. Genome Res. 23, 419–430 (2013).

71. Thibault-Sennett, S. et al. Interrogating the Functions of PRDM9 Domains in Meiosis. Genetics 209, 475–487 (2018).

72. van Belzen, I. A. E. M., Schönhuth, A., Kemmeren, P. & Hehir-Kwa, J. Y. Structural variant detection in cancer genomes: computational challenges and perspectives for precision oncology. Npj Precis. Oncol. 5, 15 (2021).

73. Creason, A. et al. A community challenge to evaluate RNA-seq, fusion detection, and isoform quantification methods for cancer discovery. Cell Syst. 12, 827–838.e5 (2021).

74. Van Twisk, D., Vincent, B. & Rubinsteyn, A. Comparing Long Read Fusion Callers using Simulated Read Data. http://biorxiv.org/lookup/doi/10.1101/2022.09.23.509226 (2022) doi:10.1101/2022.09.23.509226.

