## Supplemental Material for "A complete digital karyotype of the B-cell leukemia REH cell line resolved by long-read sequencing"

1. List of Supplemental Tables
2. Supplemental Data
3. Supplemental Figures
    - Supplemental Figure S1. TBL1XR1 deletion
    - Supplemental Figure S2. NR3C1/ARHGAP26 deletion
    - Supplemental Figure S3. BTG1 deletion
    - Supplemental Figure S4. NFATC1 deletion
    - Supplemental Figure S5. VPREB1 deletion
    - Supplemental Figure S6. Using ONT ultralong reads to call and confirm structural variants
4. Supplemental Discussion
    - Resolution of Karyotype Inconsistencies
    - Split-read analysis of der(16)t(16;21)
    - Long-read WGS outperformed for translocation discovery and resolving highly repetitive regions
    - Fusion gene splicing and discovery
    - Comparison of fusion gene caller performance
    - Instability in the CBFA2T3 region
    - The PHAX-AC007450.2 fusion
    - De-novo assembly
5. Supplemental Methods
    - G-banding
    - Automated filtering of SV candidates
    - Automated filtering of fusion gene candidates
    - Visualization
    - De-novo assembly
6. References

### 1. List of Supplemental Tables

Table S1. NCBI/SRA accession numbers

Table S2. Comparison of chromosomal features across karyotypes

Table S3. WGS depth of coverage for diploid vs. aneuploid chromosomes

Table S4. Comparison of structural variant caller performance

Table S5. Breakpoints of high-confidence fusion gene variants

Table S6. Comparison of fusion gene caller performance

### 2. Supplemental Data

Supplemental data are described and available for download at: https://zenodo.org/record/7702098. Filtering scripts are available at: https://github.com/Molmed/REH.

### 3. Supplemental Figures

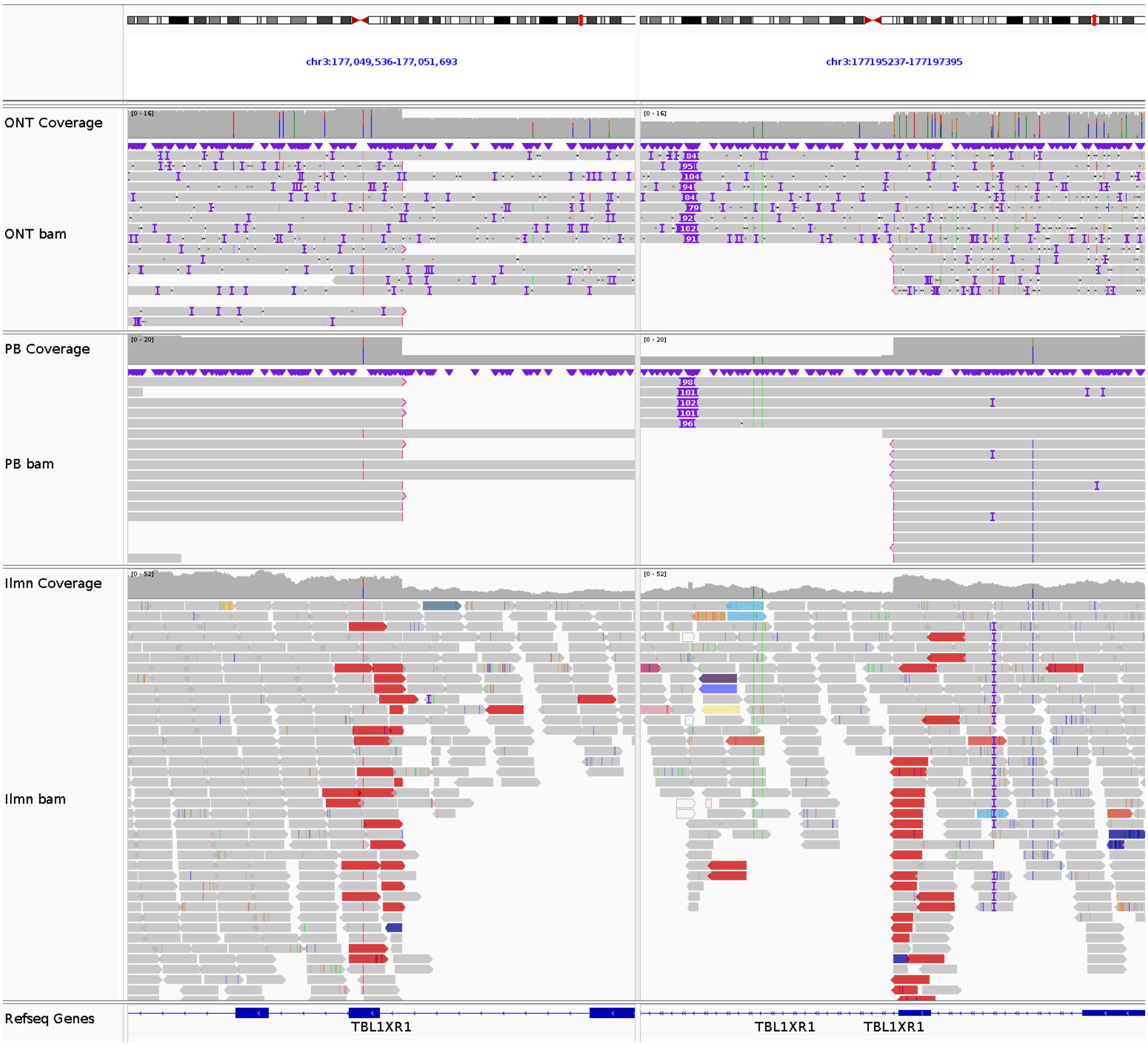


**Supplemental Figure S1. *TBL1XR1* deletion.** The breakpoints of the 146 kbp del(3)(q26.32) at chr3:177050707 and chr3:177196318, supported by 5 ONT reads, 12 PB reads and 25 Illumina reads.

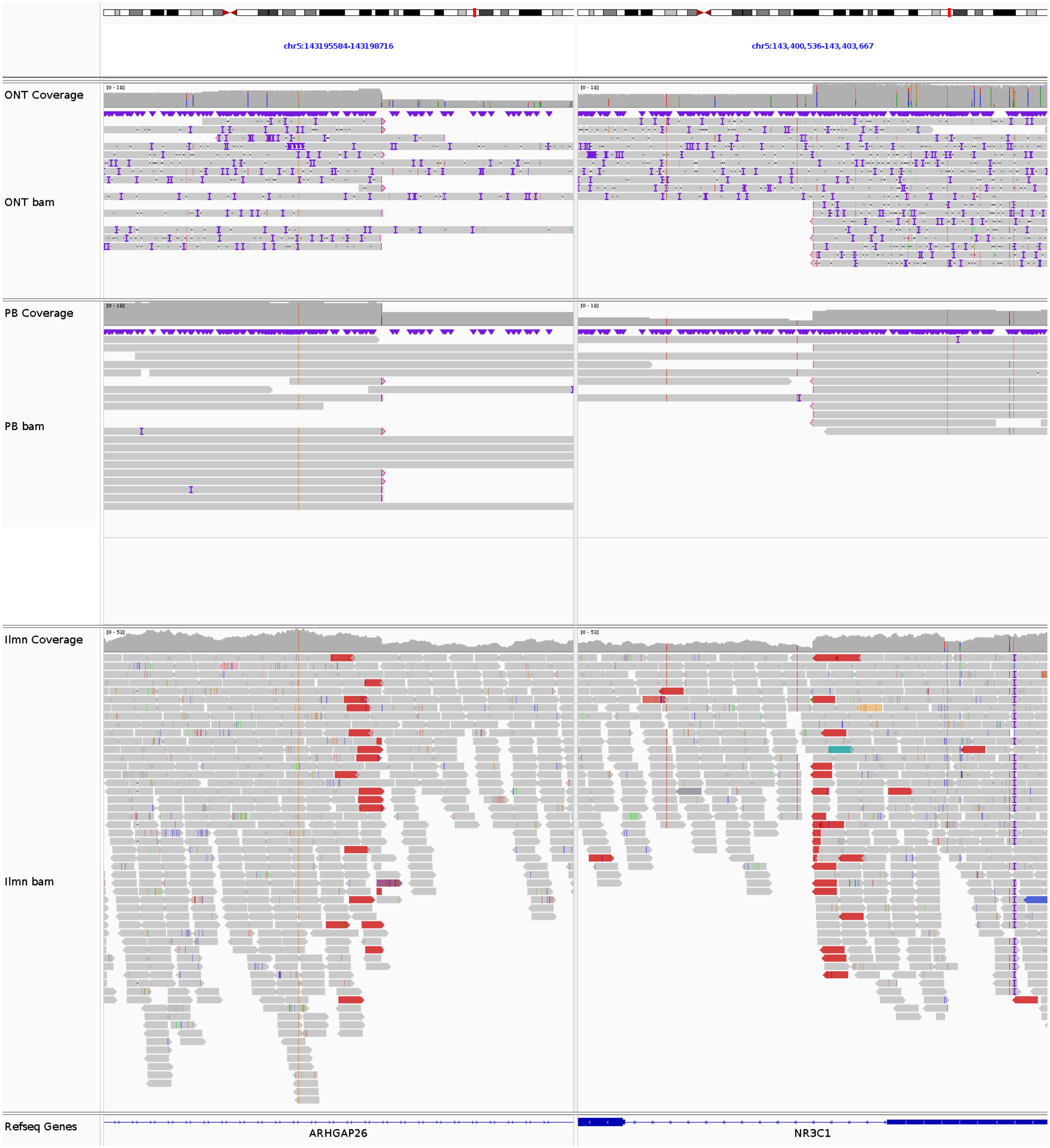


**Supplemental Figure S2.** ***NR3C1/ARHGAP26*** **deletion.** The breakpoints of the 205 kbp del(5)(q31.3) at chr5:143197445 and chr5:143402107, supported by 8 ONT reads, 7 PB reads and 19 Illumina reads.

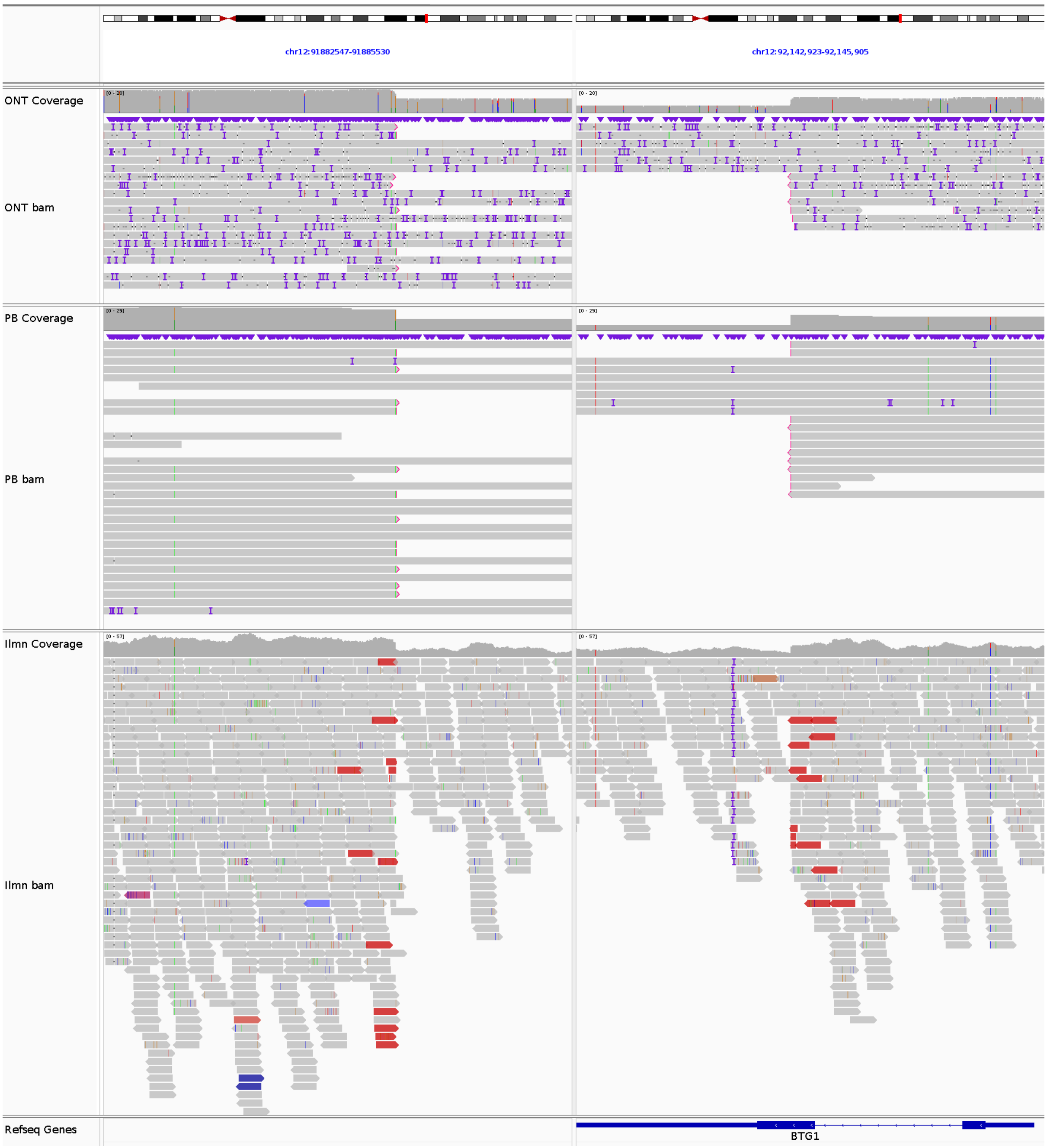


**Supplemental Figure S3. *BTG1* deletion.** The breakpoints of the 260 kbp del(12)(q21.33) at chr12:91884416 and chr12:92144292, supported by 8 ONT reads, 12 PB reads and 13 Illumina reads.

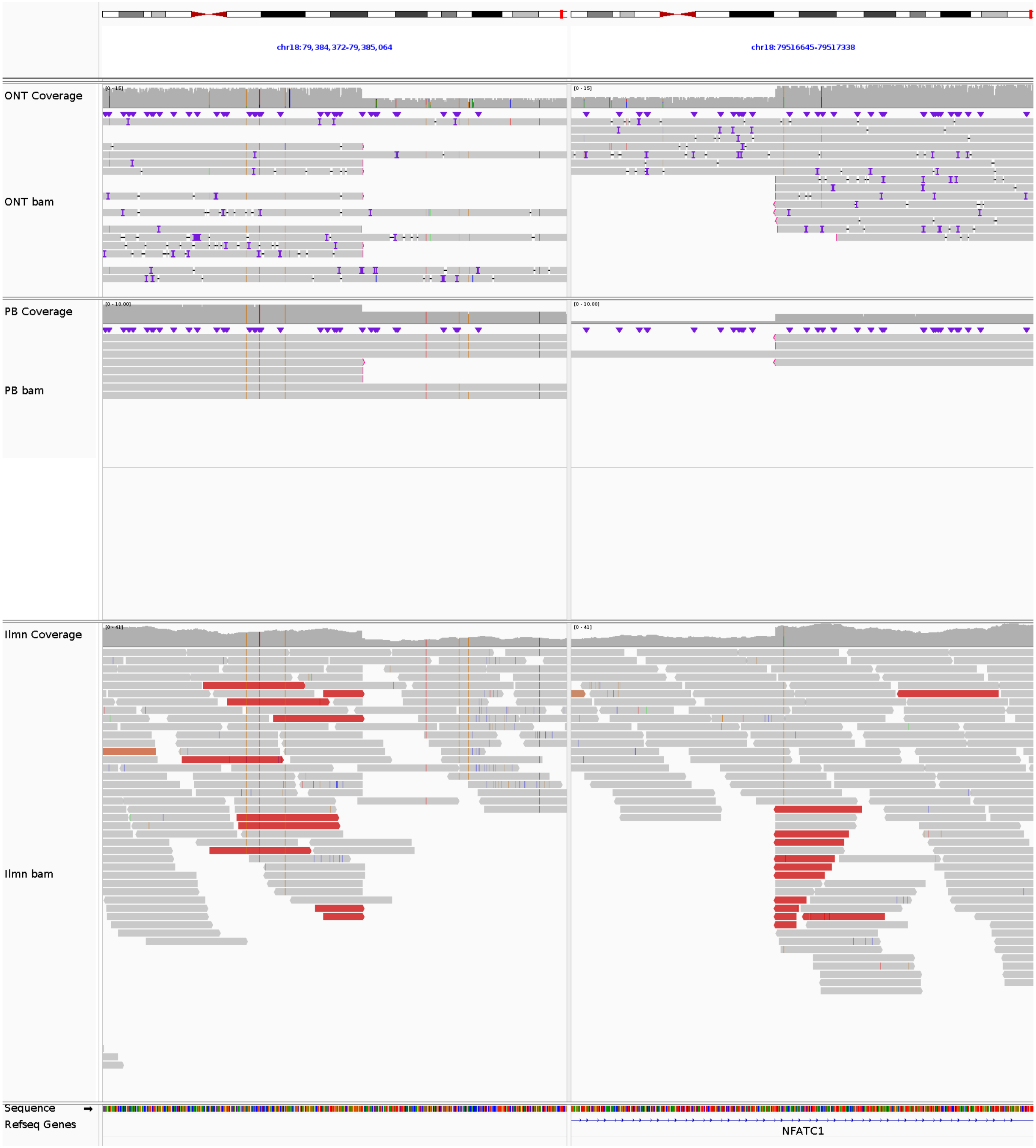


**Supplemental Figure S4.** ***NFATC1*** **deletion.** The breakpoints of the 132 kbp del(18)(q23) at chr18:79384761 and chr18:79516951, supported by 7 ONT reads, 3 PB reads and 10 Illumina reads.

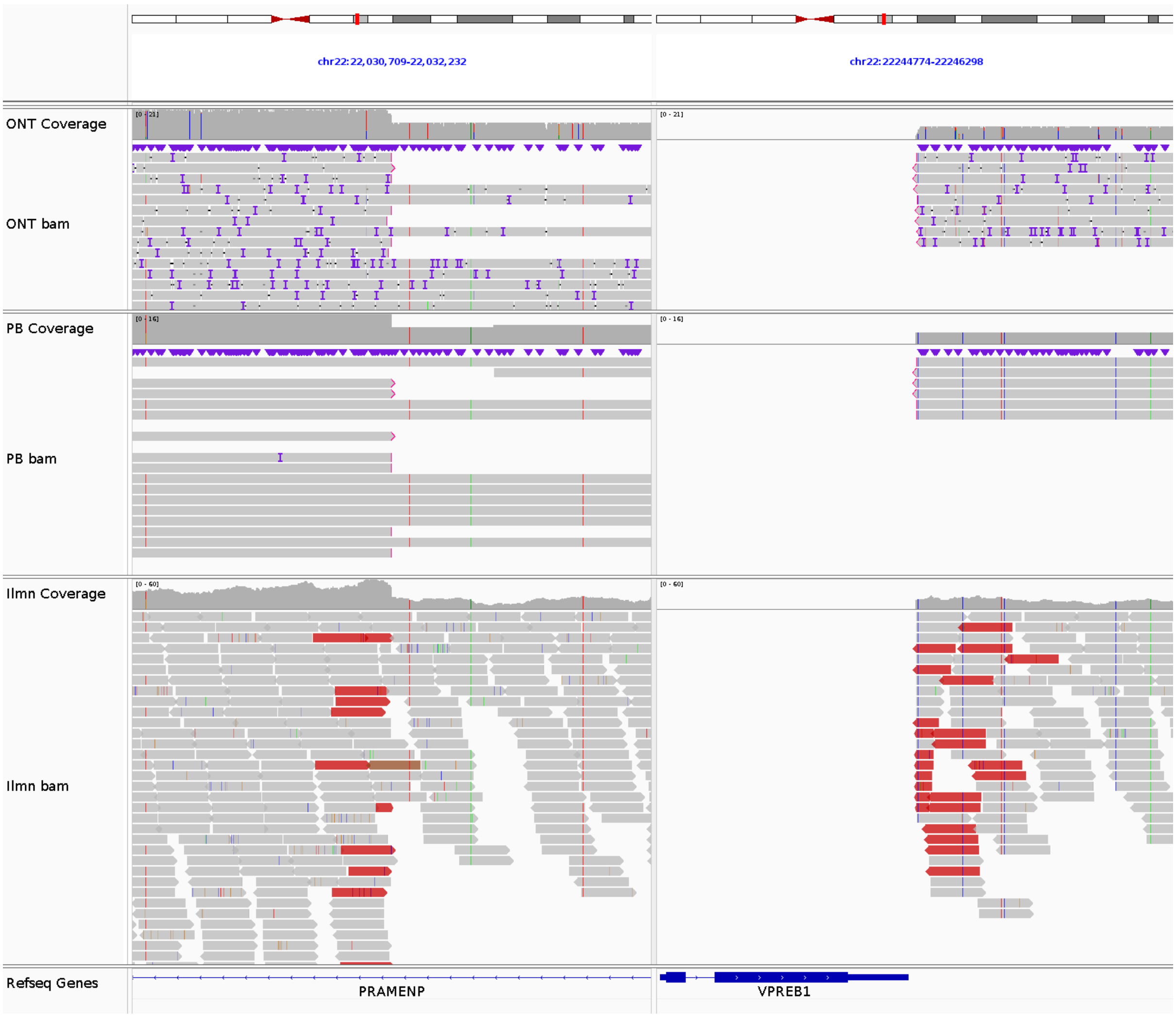


**Supplemental Figure S5. *VPREB1* deletion.** The breakpoints of the 214 kbp del(22)(q11.22) at chr22:22031472 and chr22:22245538, supported by 9 ONT reads, 6 PB reads and 29 Illumina reads.

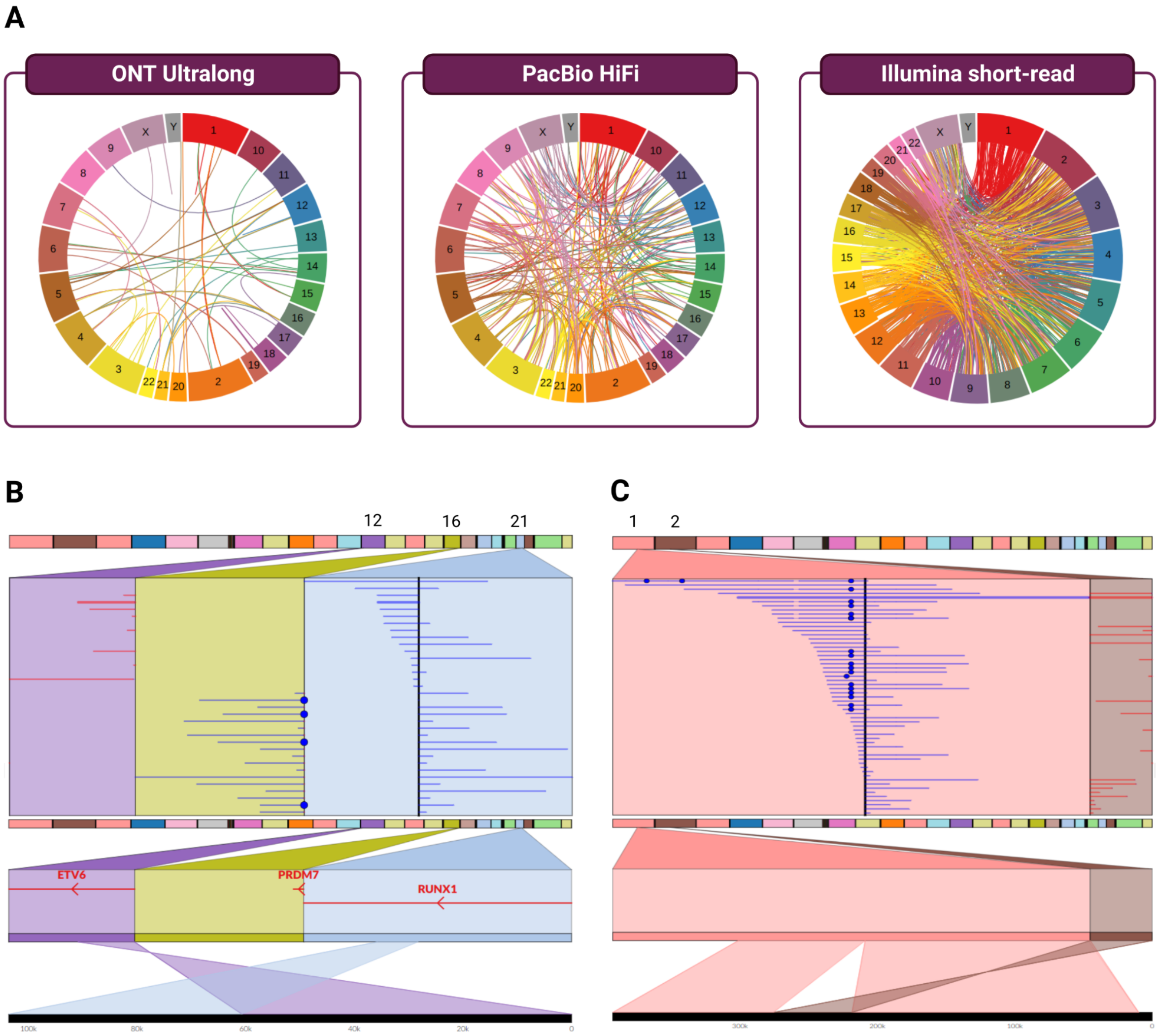


**Supplemental Figure S6. Using ONT ultralong reads to call and confirm structural variants.** a) Interchromosomal breakends detected in ONT, PacBio and Illumina short reads. The false positive rate for the called SVs was 84.7% in ONT, 96.5% in PacBio and 98.6% in the Illumina data. Visualization of the b) threeway breakpoint t(12;21;16) and c) novel t(1;2) translocation in ONT ultralong reads using Ribbon.

### 4. Supplemental Discussion

#### Resolution of Karyotype Inconsistencies

The DSMZ karyotype documents the presence of a sideline containing an extra chromosome 18 and an inversion at chromosome 5; neither of these aberrations were detected by either our G-banding nor the karyotype generated by Uphoff et al.[1], indicating that the REH cells analyzed by us did not contain this sideline (**Supplemental Table S2**).

We resolved three inconsistencies in the previously documented REH karyotypes. First, each of the three callsets resolved the breakpoints of the balanced t(5;12) to 5q23.2-23.3, with a 2.2 Mbp deletion on chr5q23 occurring between the breakpoints. The previous analog karyotypes each erroneously placed the breakpoint on chromosome 5 at q31-q32. Second, the breakpoints of the del(3) were unclear from the analog karyotypes, ranging from p13-p22 to p21.3-p24, or missing altogether. The SV callsets unanimously resolved this deletion to p14.2-p22.3. Finally, the analog karyotypes showed a discrepancy in the endpoint of the inv(12)(p13), placing it at either q22 or q23; the callsets resolved its location to q23.1.

#### Split-read analysis of der(16)t(16;21)

We confirmed the presence of two copies of the der(16)t(16;21) by analyzing the split read coverage of this breakpoint. The breakpoint was supported by 18 ONT split reads (average ONT DC chr16 = 26.4 vs 18.2 for diploid chromosomes), 19 PB split reads (average PB DC chr16 = 20.8 vs 14.6 for diploid chromosomes), and 32

Illumina discordant read pairs (average Illumina DC chr16 = 51.5 vs 34.7 for diploid chromosomes) We found a single set of breakends for this translocation, chr21:34947932 and chr16:90067326 (**Table 2; Table S3**).

#### Long-read WGS outperformed for translocation discovery and resolving highly repetitive regions

We evaluated the strengths and weaknesses of short- and long-read sequencing technologies in the context of digital karyotyping of the REH cell line. We found that Sniffles run on the ONT data had the highest sensitivity and lowest false-positive rate (FPR) of the three SV callsets. Sniffles/ONT called each of the 23 confirmed interchromosomal or large-scale SVs > 100 kb (100% sensitivity), while Sniffles/PacBio called 22 (95.65%) and TIDDIT/Illumina called 21 (91.30%). The Sniffles/ONT callset contained the smallest number of false positives (127; 84.67% FPR), which was fewer than the Sniffles/PacBio callset (605; 96.49% FPR) and TIDDIT/Illumina callset (1469; 98.59% FPR) (**Figure S6; Supplemental Table S4**).

The TIDDIT callset had a high sensitivity rate, detecting all of the confirmed SVs except for those occurring in highly repetitive regions. However, it suffered from a high false-positive rate. Of the 29 SV candidates randomly sampled from the filtered TIDDIT callset, none were visibly supported by long reads in IGV; these SV candidates were called in regions containing indels > 10bp, which were resolved in the long-read data but mismapped and misidentified as translocations or other SVs in the Illumina data.

PacBio data, which had lower average DC than ONT (14.6 PB vs 18.2 ONT), did not detect any SVs that were missed in the other two callsets; however, in both the PB and ONT datasets, Sniffles detected a t(1;2) and a 116kbp dup(1), both missed in the Illumina data and both occurring in highly repetitive regions indicated by spikes in DC (**Figure 3**).

The long-read technologies facilitated the confirmation of large-scale aberrations using split-read analysis. The visualization tool Ribbon was used to visualize reads spanning complex breakpoints such as the one between chromosomes 12, 16 and 21, and to explore how specific split reads map to the reference genome. The ultralong ONT reads in particular helped resolve the complex rearrangement t(1;2)-inv(2), where several ultralong reads spanned the entire 58kbp translocated region of chromosome 2 and its flanking regions in the derivative chromosome 1. (**Supplemental Figure S6**).

#### Fusion gene splicing and discovery

The *ETV6-RUNX1* fusion was detected in both the Illumina RNA-seq and IsoSeq datasets (five and 62 supporting reads, respectively) with support from five fusion callers. We detected two splice variants of *ETV6-RUNX1*, in line with the genomic breakpoints found in intron 5 of *ETV6* and intron 1 of *RUNX1*; however in the most prevalent *ETV6-RUNX1* transcript, exon 5 of *ETV6* was spliced to exon 2 of *RUNX1* **(Figure 4a).**

*RUNX1-PRDM7* was highly expressed, detected by all seven fusion callers, and supported by 144 Illumina reads and 24 IsoSeq reads. The genomic breakpoint in

*PRDM7* was in intron 4; five splicing variants of the *RUNX1-PRDM7* were found involving exons 5-11, with the most prevalent fusion transcript taking place between exon 1 of *RUNX1* and exon 9 of *PRDM7* (**Figure 4b**).

*PHAX-AC007450.2* and *LRP6-SLC27A6* were both detected in both the short and long-read RNA-seq datasets. In *PHAX-AC007450.2,* which was supported by nine Illumina reads and five IsoSeq reads, exon 4 of *PHAX* was fused with exon 2 of *AC007450.2*, which lies 85kb downstream of *ETV6* (**Figure 4c**). Exon 22 of *LRP6* was fused with exon 2 of *SLC27A6* (**Figure 4d**). *LRP6-SLC27A6* was supported by one short read and two long reads. However, manual inspection of the reads revealed an additional 11 low-quality split long reads that were discarded by the fusion caller and not reported in the supporting read count.

The *BTG1-LINC02404/AC090049.1* fusion was only called in the short-read RNA-seq dataset using Arriba, but by no long read callers. The resulting out-of-frame, truncated transcript was highly expressed with 120 supporting Illumina reads and nine IsoSeq reads. (**Figure 4e**).

The 205 kbp del(5)(q31.3) resulted in the antisense transcript *NR3C1-ARHGAP26*, fusing exon 1 of *NR3C1* with exon 20 of *ARHGAP26*. This fusion was only detected by one short-read fusion caller, with 10 supporting Illumina reads, but also found support in the long-read data, with five supporting IsoSeq reads. (**Figure 4f**)

Finally, the 137kb del(6)q21 resulted in the fusion gene *TRAF3IP2-REV3L*, fusing exon 8 of *TRAF3IP2* to exon 3 of *REV3L* (**Figure 4g**). This fusion was not detected by any long-read callers, but was found by three short-read fusion callers and was supported by six Illumina reads and 13 IsoSeq reads.

Splicing and read support details can be found in **Supplemental Table S5**, while fusion software support details can be found in Supplemental Data file *REH.fusions.filtered.csv*.

#### Comparison of fusion gene caller performance

The number of fusion gene candidates found by the short-read callers had a wide range: STAR-fusion found only seven, Arriba found 31, and FusionCatcher found 78; while at the higher end of the range were Squid (n = 393) and pizzly (n = 5520). The two long-read callers both returned a large number of candidates: 336 from Cupcake and 4927 from JAFFAL. The percentage of fusion gene candidates passing automated filtering were as follows: JAFFAL, 0.24%; Squid: 0.25%; pizzly, 0.36%; Cupcake: 1.79%; FusionCatcher, 7.69%; Arriba: 29.03%; and STAR-fusion: 57.14%.

The overall performance of the fusion callers also varied widely, as measured by sensitivity (percentage of the manually confirmed fusion genes detected) and FPR, calculated after discounting fusion gene candidates also found in the GM12878 RNA-seq dataset. Arriba detected each of the seven of the manually confirmed fusion genes (100.0% sensitivity, but 76.67% FPR) and STAR-fusion had the lowest FPR (50.0%) but only detected three fusion genes (42.86% sensitivity). Squid had the lowest sensitivity rate (14.29%; 1 confirmed gene fusion found), with an FPR of 99.65%. The remaining tools each called three of the confirmed fusion genes

(42.86% sensitivity rate) and had an FPR between 94.23-99.94%. (**Supplemental Table S6**).

#### Instability in the *CBFA2T3* region

The fusion callers suggested several other fusion events between *RUNX1* and genes proximal to the genomic breakpoint on 16q24.3. Notably, the *CBFA2T3* gene was suggested as a fusion partner by both long-read callers. Manual inspection did not confirm the fusions involving *CBFA2T3*; however, there were at least four large indels of 90-800bp within this region, as well as a number of smaller indels (< 50bp), suggesting a genomic instability in the region disrupting the gene's structure.

We found evidence of genomic instability throughout chr16q24.3, including the region encompassing *CBFA2T3*, 1mbp downstream of *PRDM7*. Aberrations involving chr16q24.3 are frequent in ALL as well as in acute myeloid leukemia (AML), where one of the well-established subtypes is defined by t(16;21) *RUNX1-CBFA2T3* [2]. REH has abnormally high expression of both *CBFA2T3* and *RUNX1* [3] and the RUNX1 fusion activates a topological disruption of multiple genes in this region that promotes BCP-ALL proliferation [4].

#### The *PHAX-AC007450.2* fusion

One of the two fusion genes arising from the balanced t(5;12) is PHAX-AC007450.2. AC007450.2 refers to the Bacterial Artificial Chromosome (BAC) clone RP11-434C1 (https://www.ncbi.nlm.nih.gov/nuccore/AC007450.1), which is 85 kb downstream, and on the opposite strand of, ETV6 [57,74]. PHAX-AC007450.2 has not been reported in ALL cases previously; however, PHAX, a protein-coding gene involved in the nuclear export of small nuclear RNA [75], appears in a single PHAX-IGH fusion that was recently reported in a ETV6-RUNX1-like case [76].

#### De-novo assembly

We created three de-novo assemblies using hifiasm on PacBio Hifi reads; Flye on ONT reads with Medaka polishing; and Flye on ONT reads followed by polishing with PacBio HiFi reads using racon. The hifiasm assembly generated 2580 contigs, N50 of 3.5Mbp and L50 of 255. The ONT-Medaka assembly generated 2060 contigs, N50 of 58Mbp and L50 of 20. The ONT-racon assembly generated 1712 contigs, N50 of 58.2Mbp and L50 of 19. We provide these assemblies as a supplement to the present study.

There are two methods for resolving genomes: alignment to a reference genome, as we have used in the present study, and de-novo assembly. Within an oncological context, the alignment approach may work better, as de-novo assembly may struggle to construct accurate contigs due to the complexity and heterozygosity of cancer genomes[5]. On the other hand, a known challenge in detecting and interpreting SVs is differentiating which are somatic and specific to pathological tissue, as relying on reference genomes makes it difficult to take into account a patient's normal genomic variation. Where matched tumor-normal samples are available, sequencing and de-novo assembly of both samples, followed by a comparative analysis, can be used for accurate, personalized SV detection[6]. In the present study, we briefly explored the possibilities of de-novo assembly, and the assemblies we generated are herein made available for further investigation of the REH cell line.

### 5. Supplemental Methods

#### G-banding

The cells from the cell culture were harvested following standard cytogenetic methods (KaryoMAX Colcemid Solution in HBSS, Gibco™), and chromosome preparations were stained by G-banding trypsin-Giemsa standard procedures (0.25% Trypsin, Gibco™, Wright stain, Sigma Aldrich, Giemsa's stain, VWR Chemicals and Buffer tablets pH 6.8, Merck). Twenty-five metaphases were analyzed using a Carl Zeiss scanner microscope and Metafer and Ikaros software from MetaSystems. The karyotype was carried out according to the International System for Human Cytogenetic Nomenclature (ISCN) 2020.

#### Automated filtering of SV candidates

For the evaluation of large-scale variants and translocations, the callsets were pre-filtered, retaining SV candidates with either length > 100000, or of a breakend type and mapping to two different chromosomes; discarded candidates were excluded from accuracy statistics. The remaining candidates were further filtered, requiring them to meet all of the following criteria: 1) passes all quality filters set by the SV calling software 2) a minimum of five supporting reads, with the number of supporting reads > 20% of the average depth of coverage (DC) for the dataset 3) position coverage is no greater than 150% of the average DC for the dataset.

Of the remaining candidates, the following were selected for manual inspection: 1) all candidates with support in at least one Sniffles callset 2) TIDDIT candidates with exactly 15 supporting reads, providing a pseudo-random sampling from this callset.

#### Automated filtering of fusion gene candidates

We filtered the fusion gene candidates from the rnafusion, JAFFAL and Cupcake outputs. All candidate fusion events called by rnafusion in the GM12878 cell line were removed and excluded from accuracy statistics. The remaining candidate fusion genes were required to have different partner genes and to fulfill any of the following criteria: 1) contains one or more known ALL-associated genes [5], and is supported by at least 5 reads 2) called by at least one short-read fusion detection tool and one long-read fusion detection tool 3) called by at least three short-read fusion detection tools, or 4) supported by at least 10 long reads.

#### Visualization

We ran Copycat v9e21f79 (https://github.com/marianattestad/copycat) on the short- and long-read BAM files in order to bin the read coverage in preparation of visualization. Visualization was performed with SplitThreader [6], Ribbon [7] and IGV. We used SplitThreader to visualize interchromosomal SVs across the Illumina, PacBio and ONT callsets, while Ribbon was used for inspection of intrachromosomal features and split-read analysis. Circos plots were drawn using Circa v1.2.3 (https://omgenomics.com/circa/).

#### De-novo assembly

We created three de-novo assemblies: one using PacBio CCS reads and hifiasm[8] v0.16.1-r375 with no pre-processing or polishing; and two different assemblies using the ONT reads. The quality of the assemblies was assessed with QUAST[9] v5.2.0.

Adapter sequences were trimmed from the ONT data prior to assembly using Porechop v0.2.4. We additionally trimmed the first 50bp from all reads and removed all reads shorter than 500bp using NanoFilt[10]. Both assemblies were perfomed using Flye[11] v2.9.1-b1780. For this first ONT assembly approach (ONT-Medaka), we performed polishing using Medaka (v1.7.0, https://github.com/nanoporetech/medaka), which uses neural networks and is trained on specific configurations of ONT pore type, instrument, and basecaller. This assembly was polished using medaka model r941_prom_hac_g507. In the second approach (ONT-Racon), we performed polishing using Racon (v1.5.0, https://github.com/lbcb-sci/racon), a graph-based polishing tool that enables error correcting with PacBio HiFi reads. In preparation, we mapped the HiFi reads to the draft assembly using minimap2 v2.24-r1122 with the "map-hifi" option enabled. We then used the resulting mapping together with the HiFi reads as input to racon.

### 6. References


1. Uphoff, C. C. *et al.* Occurrence of TEL-AML1 fusion resulting from (12;21) translocation in human early B-lineage leukemia cell lines. *Leukemia* **11**, 441–447 (1997).
2. Noort, S. *et al.* Prognostic impact of t(16;21)(p11;q22) and t(16;21)(q24;q22) in pediatric AML: a retrospective study by the I-BFM Study Group. *Blood* **132**, 1584–1592 (2018).
3. Rouillard, A. D. *et al.* The harmonizome: a collection of processed datasets gathered to serve and mine knowledge about genes and proteins. *Database* **2016**, baw100 (2016).
4. Jakobczyk, H. *et al.* Reduction of RUNX1 transcription factor activity by a CBFA2T3-mimicking peptide: application to B cell precursor acute lymphoblastic leukemia. *J. Hematol. Oncol.J Hematol Oncol* **14**, 47 (2021).
5. Marincevic-Zuniga, Y. *et al.* Transcriptome sequencing in pediatric acute lymphoblastic leukemia identifies fusion genes associated with distinct DNA methylation profiles. *J. Hematol. Oncol.J Hematol Oncol* **10**, 148 (2017).
6. Nattestad, M., Alford, M. C., Sedlazeck, F. J. & Schatz, M. C. *SplitThreader: Exploration and analysis of rearrangements in cancer genomes*. http://biorxiv.org/lookup/doi/10.1101/087981 (2016) doi:10.1101/087981.
7. Nattestad, M., Aboukhalil, R., Chin, C.-S. & Schatz, M. C. Ribbon: intuitive visualization for complex genomic variation. *Bioinformatics* **37**, 413–415 (2021).
8. Cheng, H., Concepcion, G. T., Feng, X., Zhang, H. & Li, H. Haplotype-resolved de novo assembly using phased assembly graphs with hifiasm. *Nat. Methods* **18**, 170–175 (2021).
9. Mikheenko, A., Prjibelski, A., Saveliev, V., Antipov, D. & Gurevich, A. Versatile genome assembly evaluation with QUAST-LG. *Bioinformatics* **34**, i142–i150 (2018).
10. De Coster, W., D'Hert, S., Schultz, D. T., Cruts, M. & Van Broeckhoven, C. NanoPack: visualizing and processing long-read sequencing data. *Bioinformatics* **34**, 2666–2669 (2018).
11. Kolmogorov, M., Yuan, J., Lin, Y. & Pevzner, P. A. Assembly of long, error-prone reads using repeat graphs. *Nat. Biotechnol.* **37**, 540–546 (2019).
